# Universal modulation of liquid-liquid phase separation by macromolecules

**DOI:** 10.64898/2026.09.03.749048

**Authors:** Yunxiao Kan, Jie Lin

## Abstract

Liquid-liquid phase separation (LLPS) widely organizes the cellular interior, yet how third-party macromolecules modulate LLPS remains a puzzle. While one would expect that macromolecules can either promote or suppress phase separation, empirical observations show that a small dosage of macromolecules virtually always promotes phase separation. To resolve this paradox, we study the linear response of phase separation to macromolecule addition. We prove that for macromolecules significantly larger than the solvent molecule, their addition essentially always promotes phase separation, regardless of how they interact with the scaffold protein. Crucially, in the biologically relevant scenario of dilute condensates, the linear responses, including the saturation volume fraction, scale proportionally to [1 − *v*_2_(1 + *χ*_12_)^2^], where *v*_2_ is the macromolecule size and *χ*_12_ is the scaffold-macromolecule interaction parameter. Importantly, this universal behavior extends to multicomponent solutions in which the effective interaction parameter is composition-dependent. Our work shifts macromolecular modulation of LLPS from empirical observation to predictable design.

## Introduction

Biomolecular condensates, often formed through liquid-liquid phase separation (LLPS), have emerged as a fundamental mechanism for spatial organization inside cells [1–5]. These membraneless organelles compartmentalize proteins and nucleic acids to regulate diverse biological processes, including signal transduction, stress response, and gene expression [6–8]. While the cellular interior is a crowded multicomponent environment [9], it is well established that the formation of biomolecular condensates is often driven by a scaffold protein (or a few of them), which is both necessary for condensate formation and capable of phase separating into droplets on its own [10–15].

Thus, a critical question naturally arises: how does a third-party macromolecule modulate the phase separation of the scaffold protein? Intriguingly, experiments have found that crowding agents, e.g., Ficoll, dextran, and polyethylene glycol (PEG), virtually always promote phase separation [9, 10, 16–18]. For example, droplets emerge upon the addition of a macromolecule to an originally homogeneous solution, indicating that adding these crowding agents reduces the scaffold protein’s saturation volume fraction, i.e., the minimum volume fraction required to trigger phase separation.

Meanwhile, RNA also almost always promotes phase separation of the scaffold protein [10, 12, 13, 15, 16, 19– 23], as long as the RNA concentration is not so high that reentrant phenomena occur [3]. These observations appear counterintuitive, as RNAs can presumably interact with the scaffold protein either repulsively or attractively. Why does the addition of a macromolecule almost always promote phase separation of the scaffold protein? Can we predict whether macromolecule addition promotes or suppresses phase separation? Despite a wealth of experimental data, a unified theoretical framework that addresses these questions remains elusive. In this work, we address this long-standing issue and focus on the linear-response regime, where the saturation volume fraction of the scaffold protein is linearly dependent on the dosage of the added macromolecule.

Based on a simple model of three-component solution [2, 24–26], we reveal an “emergent simplicity” in the biologically relevant scenario where the volume fraction of the scaffold protein is low inside condensates: the linear response of the protein saturation volume fraction (*ϕ*_1*a*_) against the added macromolecule volume fraction 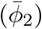 becomes 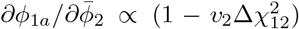. Here, *v*_2_ is the macromolecule size relative to the solvent molecule (i.e., water), and Δ*χ*_12_ = *χ*_12_ + 1 where *χ*_12_ is the Flory interaction parameter between the macromolecule and the scaffold protein. Our theory predicts that, for large macromolecules with *v*_2_ ≫ 1, regardless of whether they interact with the scaffold protein attractively or repulsively, their addition virtually always promotes the phase separation of the scaffold protein by lowering its saturation volume fraction (Fig. 1a). Meanwhile, they also increase the dense-phase volume fraction and reduce the total size of preexisting condensates (Fig. 1b).

**FIG 1.**
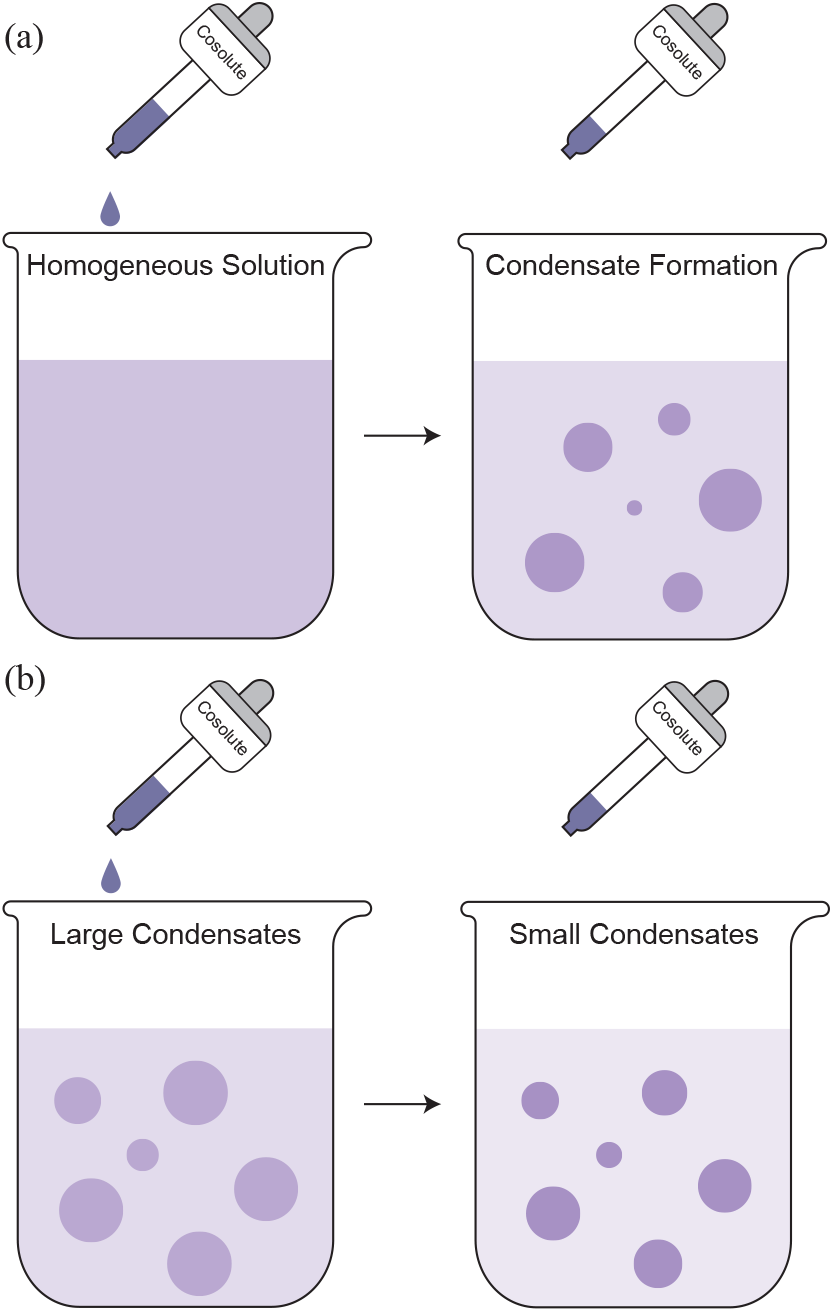
Two critical effects of macromolecule addition.(a) Macromolecule addition virtually always promotes phase separation in the low-dosage limit by lowering the scaffold protein’s saturation volume fraction; e.g., adding a macromolecule to a homogeneous solution induces condensate formation. (b) Macromolecule addition also virtually always reduces the volume ratio of preexisting condensates, i.e., leading to smaller but more concentrated condensates.

Furthermore, we extend our theory to multicomponent solutions and demonstrate that our conclusions are valid for all biomolecule species in a multicomponent solution as long as *χ*_12_ is replaced by *χ*_eff_, the effective interaction parameter between the added macromolecule and all the preexisting biomolecules. Our theory rationalizes the widely observed promoting effects of macromolecules on phase separation and provides a predictive framework for understanding how cells might tune their internal composition to regulate condensate formation and dissolution.

## Model

We consider a three-component solution containing the scaffold protein, the macromolecule, and the solvent (we will extend our theory to multicomponent solutions with more than three components later). We model the free energy density using the Flory-Huggins theory [24, 25]

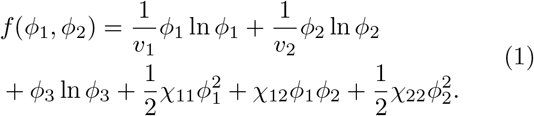

Here, *ϕ*_1_, *ϕ*_2_, and *ϕ*_3_ = 1 − *ϕ*_1_ − *ϕ*_2_ are the volume fractions of the protein, macromolecule, and the solvent, respectively. We set the volume of the solvent molecule, *v*_0_, as the volume unit, and the dimensionless volumes of the protein and the macromolecule are *v*_1_ and *v*_2_, respectively. The unit of free energy density is *k*_*B*_*T/v*_0_, where *k*_*B*_ is the Boltzmann constant and *T* is the temperature. *χ*_11_, and *χ*_12_, *χ*_22_ are the Flory interaction parameters.

Considering a protein solution with a dilute (a) and dense (b) phase, we introduce the volume ratios, *f*_*a*_ and *f*_*b*_, as the ratios of their volumes to the total solution volume, which satisfy *f*_*a*_ +*f*_*b*_ = 1. At equilibrium, the chemical potentials of the protein and macromolecule, denoted by *µ*_1_ and *µ*_2_, respectively, and the osmotic pressure Π, must be equal between the two phases. *ϕ*_1*a*_, *ϕ*_1*b*_, *ϕ*_2*a*_, *ϕ*_2*b*_, *f*_*a*_ are uniquely determined by the following equilibrium and conservation conditions: (1) *µ*_1*a*_ = *µ*_1*b*_, (2) *µ*_2*a*_ = *µ*_2*b*_, (3) Π_*a*_ = Π_*b*_, (4) 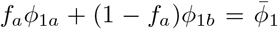, and (5) 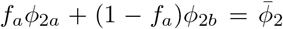. Here, *µ*_*i*_ = ∂*f/*∂*ϕ*_*i*_ where *i* = 1, 2, and 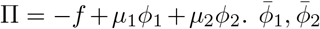 are the average volume fractions of the two solutes in the whole solution [27]. The linear responses of *ϕ*_1*a*_, *ϕ*_1*b*_, *ϕ*_2*a*_, *ϕ*_2*b*_, and *f*_*a*_ to a small change in 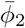 can be found by solving the following linear equations:

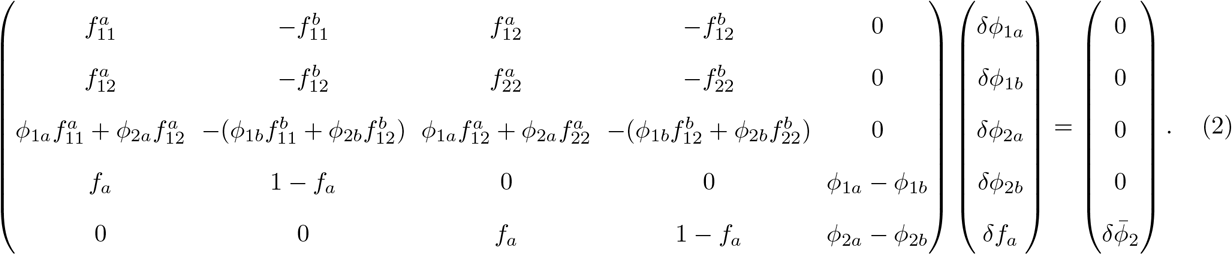

Here, 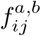 is defined as ∂^2^*f/*∂*ϕ*_*i*_∂*ϕ*_*j*_ and the upper index represents the dilute (a) or dense (b) phases.

### Effects of macromolecule addition on the saturation volume fraction

We study how the macromolecule addition affects the saturation volume fraction of the scaffold protein, i.e., *ϕ*_1*a*_, a biologically critical variable. We remark that directly solving Eq. (2) is cumbersome. Applying Cramer’s rule and Laplacian theorem to Eq. (2) (Supplemental Material (SM) section D3), we obtain the linear response of *ϕ*_1*a*_ analytically, which has not been reported before to the best of our knowledge:

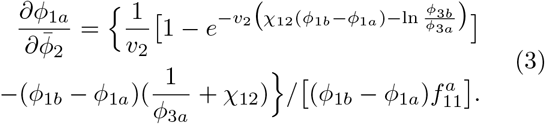

In this work, 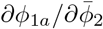 is computed at the limits 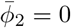 and *f*_*a*_ = 1 unless otherwise mentioned. In Eq. (3), *ϕ*_1*a*_ *ϕ*_1*b*_ are the protein volume fractions of the dilute and dense phase before macromolecule addition, with 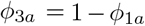 and *ϕ*_3*b*_ = 1 − *ϕ*_1*b*_. We remark that taking *f*_*a*_ = 1 in Eq. (3) not only simplifies the derivation but also lets 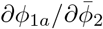 have an intuitive interpretation: the local slope of the phase boundary at 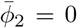 (−tan *θ* in Fig. 2b inset; see a detailed derivation in SM section C4). We confirm Eq. (3) by numerically computing the protein saturation volume fractions (Fig. S1).

**FIG 2.**
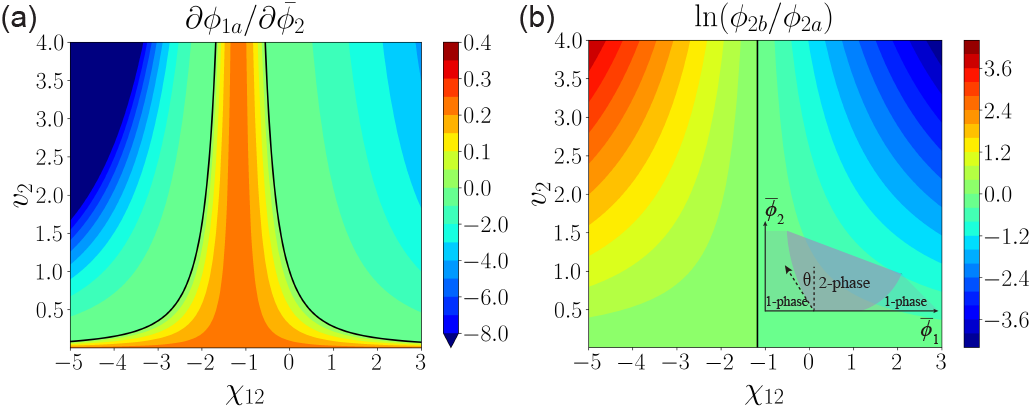
Linear response of the saturation volume fraction to macromolecule addition. (a) Heat map of 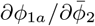 as a function of *χ*_12_ and *v*_2_ according to Eq. (3). To apply Eq. (3), we first compute *ϕ*_1*a*_ and *ϕ*_1*b*_ for the two-component system using the numerical method from Ref. [28]. The black solid curves represent the zero-level curves along which 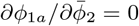. (b) Heat map of ln(*ϕ*_2*b*_*/ϕ*_2*a*_) in the linear response regime where *ϕ*_2*a*_ and *ϕ*_2*b*_ are the volume fractions of the macromolecule in the dilute and dense phases, respectively. The black line represent the curve along which *ϕ*_2*b*_ = *ϕ*_2*a*_. The inset shows the geometrical meaning of 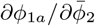 at *f*_*a*_ = 1. In this figure, *v*_1_ = 100, *χ*_11_ = −1.3.

We highlight a special point in the parameter space (*v*_2_ = 1, *χ*_12_ = 0) where the macromolecule has the same size as the solvent, and the interaction strength between protein and macromolecule is zero. At this point, 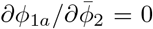 according to Eq. (3), as the protein cannot distinguish the molecule from the solvent anymore. In fact, the protein saturation volume fraction is constant for any 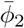 if *v*_2_ = 1, *χ*_12_ = *χ*_22_ = 0 (SM section C2).

Interestingly, we identify two zero-level curves, 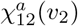 and 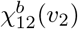 along which 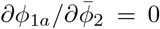 (see the black lines in Fig. 2a). Macromolecules that have either a strong attractive or a strong repulsive interaction with the protein can promote phase separation. To understand this observation, we compute the partition coefficient (PC), i.e., the ratio of the macromolecule volume fraction in the dense phase to that in the dilute phase, *P* = *ϕ*_2*b*_*/ϕ*_2*a*_. In the linear response regime, 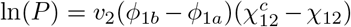 (SM section D1). Here, we introduce the critical protein-macromolecule interaction parameter, 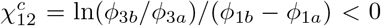. Intuitively, for large positive *χ*_12_, the macromolecule is enriched in the dilute phase (Fig. 2b); the repulsion between protein and macromolecule facilitates protein accumulation in the dense phase and therefore lowers its saturation volume fraction. For very negative *χ*_12_, the macromolecule is enriched in the dense phase; the attraction between protein and macromolecule helps recruit protein into the dense phase and therefore also lowers its saturation volume fraction.

Our theory agrees with the three archetypical classes of macromolecular regulators proposed in Ref. [29]. Given a fixed *v*_2_, as *χ*_12_ decreases from a positive to a negative value (Fig. 2a), the nature of the added macro-molecule switches from volume-exclusion promoters (low PC) to weak-attraction suppressors (moderate PC), and finally to strong-attraction promoters (high PC). In what follows, we show that, in a biologically relevant scenario, the linear responses of the scaffold protein’s phase-separation properties to the addition of macromolecules exhibit emergent simplicity.

### Emergent simplicity in the biologically relevant scenario

We note that the volume fraction of the scaffold protein inside biomolecular condensates is typically low, varying from approximately 0.01 to 0.2 (in the meantime, the saturation volume fraction is often several orders of magnitude smaller than the dense-phase volume fraction) [30, 31]. Therefore, in this biologically relevant scenario, one can reasonably approximate the volume fraction of the dense phase as being much smaller than one. Meanwhile, it is known that such condensates must be composed of a long polymer, along with the condition 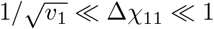 where Δ*χ*_11_ = − 1 − *χ*_11_ such that the volume fractions of the dilute and dense phases satisfy 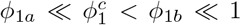 [27, 32, 33]. Intriguingly, in the biologically relevant scenario, the linear response in the protein saturation volume fraction becomes (SM section F):

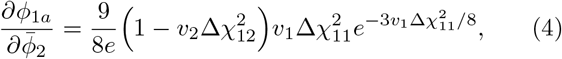

where Δ*χ*_12_ = *χ*_12_ + 1 with the zero-level curves of 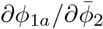 simplified as

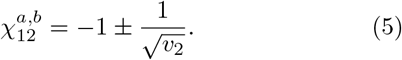

Eq. (5) predicts that large macromolecules (*v*_2_ ≫ 1) virtually always promote phase separation by decreasing the saturation volume fraction regardless of how it interacts with the scaffold protein. Furthermore, the partition coefficient is also simplified as ln *P* = −*ϕ*_1*b*_*v*_2_Δ*χ*_12_ where *ϕ*_1*b*_ = 3*/*2Δ*χ*_11_ (Fig. S3).

We next study how macromolecule affects a solution that has undergone phase separation with 0 *< f*_*a*_ *<* 1 and derive the linear responses of the dense-phase volume fraction and volume ratio of preexisting condensates in the biologically relevant scenario (SM section F and G):

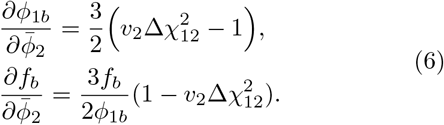

Intriguingly, the changes in *ϕ*_1*b*_ and *f*_*b*_ share the same zero-level curves as the saturation volume fraction. Interestingly, if a macromolecule promotes phase-separation of the scaffold protein by lowering its saturation volume fraction, it will also reduce the volume ratio of preexisting condensates in the same protein solution that already has two phases. This is because the reduction of condensate size is accompanied by an increase in *ϕ*_1*b*_ so that the total protein mass is still conserved (note that the majority of protein mass is in condensates). We numerically test Eqs. (5, 6) and obtain nice agreements (Fig. 3).

**FIG 3.**
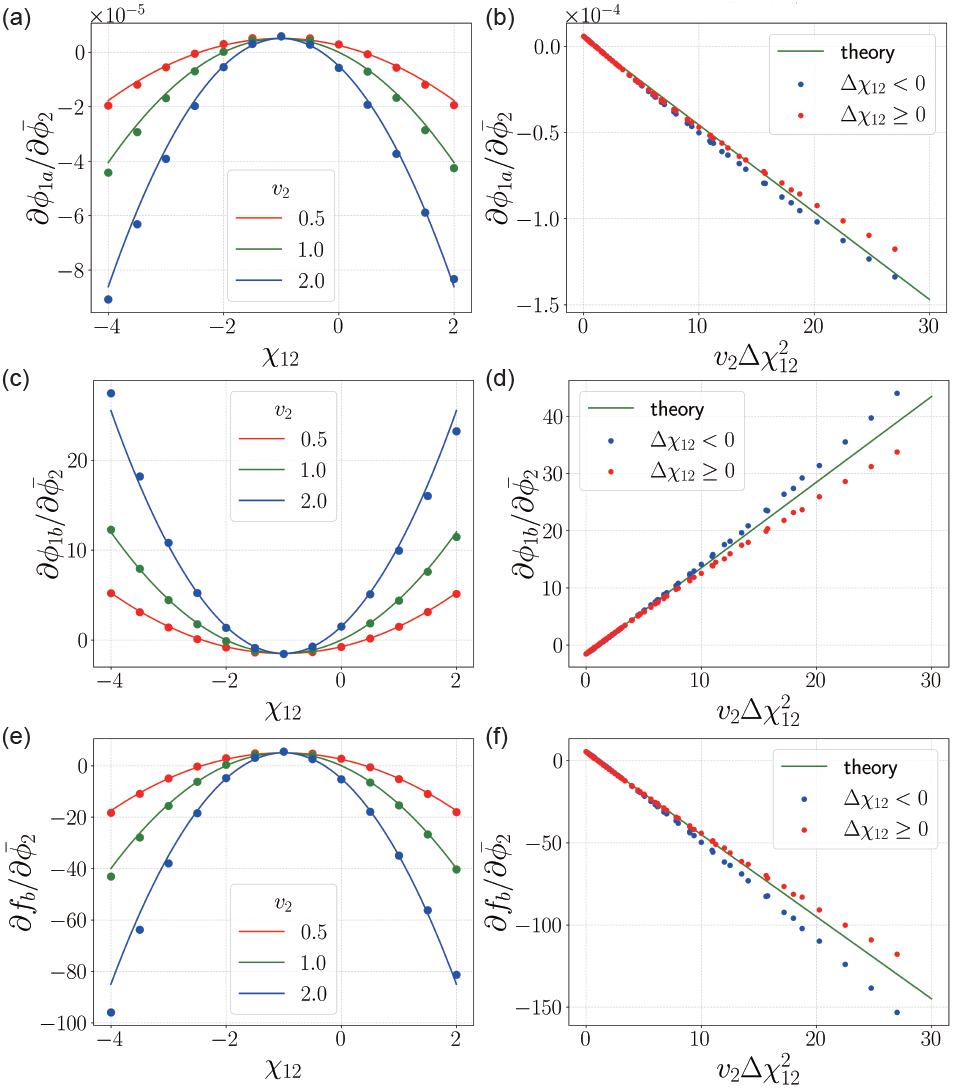
Linear responses of the protein solution to macromolecule addition. (a) The linear response of the saturation volume fraction of the scaffold protein. The data points are from numerical calculations, and the solid lines are theoretical predictions. (b) The same data as (a), but the x-axis is rescaled as 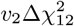. Data for different *v*_2_ collapse onto the same master curve. (c, d) The same as (a, b) but for the dense-phase volume fraction of preexisting condensates. (e, f) The same as (a, b) but for the volume ratio of preexisting condensates. In this figure, *v*_1_ = 10^5^, *χ*_11_ = − 1.02, the data points are computed from two states with 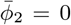 and 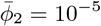 in (a, b) and *f*_*a*_ = 0.9 in (c-f).

### Multicomponent solutions

We generalize our model to N+2-component solutions with two phases: the number of preexisting biomolecule species is N before macromolecule addition, where *N >* 1. Importantly, our theory regarding the volume fractions of preexisting biomolecules can be generalized as (SM section H1):

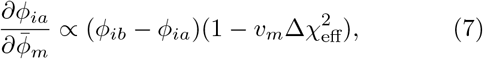

where 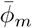 is the average volume fraction of the added macromolecule, *v*_*m*_ is the macromolecule size, and the effective interaction parameter 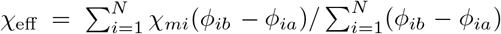. In deriving Eq. (7), we use the condition *ϕ*_*ia*_ ≪ 1, *ϕ*_*ib*_ ≪ 1 and *f*_*a*_ = 1. Note that the definition of dilute or dense phase is arbitrary in this case; instead, we call phase a the major phase and phase b the minor phase based on their volume ratio.

Strikingly, Eq. (7) predicts that the linear responses of all the biomolecule species share the same zero-level curves with the signs depending on whether they are enriched in the major or minor phase. To verify Eq. (7), we numerically generate an N+1-component solution with two phases, add a small amount of macromolecule, and track the resulting changes in *ϕ*_*ia*_. Indeed, all the N biomolecule species share the same predicted boundary separating the Δ*ϕ*_*ia*_ *<* 0 and Δ*ϕ*_*ia*_ *>* 0 regions.

## Discussion

In this work, we study the modulation of phase separation by macromolecules in the linear-response regime. Our theory explains empirical observations from simulations of the patchy particle model [34] and experiments on the SH3_5_-PRM_5_ system [29]. We also uncover the emergent simplicity in the biologically relevant scenarios. In particular, the linear responses only depend on *v*_2_(1 + *χ*_12_)^2^ for a 3-component solution.

Furthermore, we generalize our results to multicomponent solutions. Importantly, our conclusions regarding the signs of the linear responses of the major-phase volume fractions are robust and only depend on the parameter *v*_*m*_(1 + *χ*_eff_)^2^, where *χ*_eff_ is the effective interaction parameter between the macromolecule and all the preexisting biomolecules, which is composition-dependent. Our theory also agrees with the experiments of Ref. [29], where the authors found that the nature of the regulatory macromolecule, lysozyme, changes from a weak-attraction suppressor in a 1:1 SH3_5_-PRM_5_ solution to a strong-attraction promoter in a 2:1 SH3_5_-PRM_5_ solution. We also extend Eq. (7) to *f*_*a*_ *<* 1 for solutions that consist of one major scaffold protein and several minor components (SM section H2).

While we focus primarily on linear responses in this work, nonlinear effects are also important. For example, RNA can switch from a promoter to a suppressor of phase separation as its concentration increases, a phenomenon known as reentrant behavior [3, 29]. Our theory predicts that if cells produce large macromolecules to dissolve condensates, they must produce enough of these regulatory macromolecules to move beyond linear response. Finally, we note that the Flory-Huggins model of the free-energy density is intrinsically a mean-field model, although our framework applies to any free-energy form. Given the great success of the Flory-Huggins model [2, 26], our theory should capture the universal features of macromolecular modulation on phase separation. Indeed, by extending our theory to a non-mean-field version [27], we find that the zero-level curves remain the same (SM section I).

**FIG 4.**
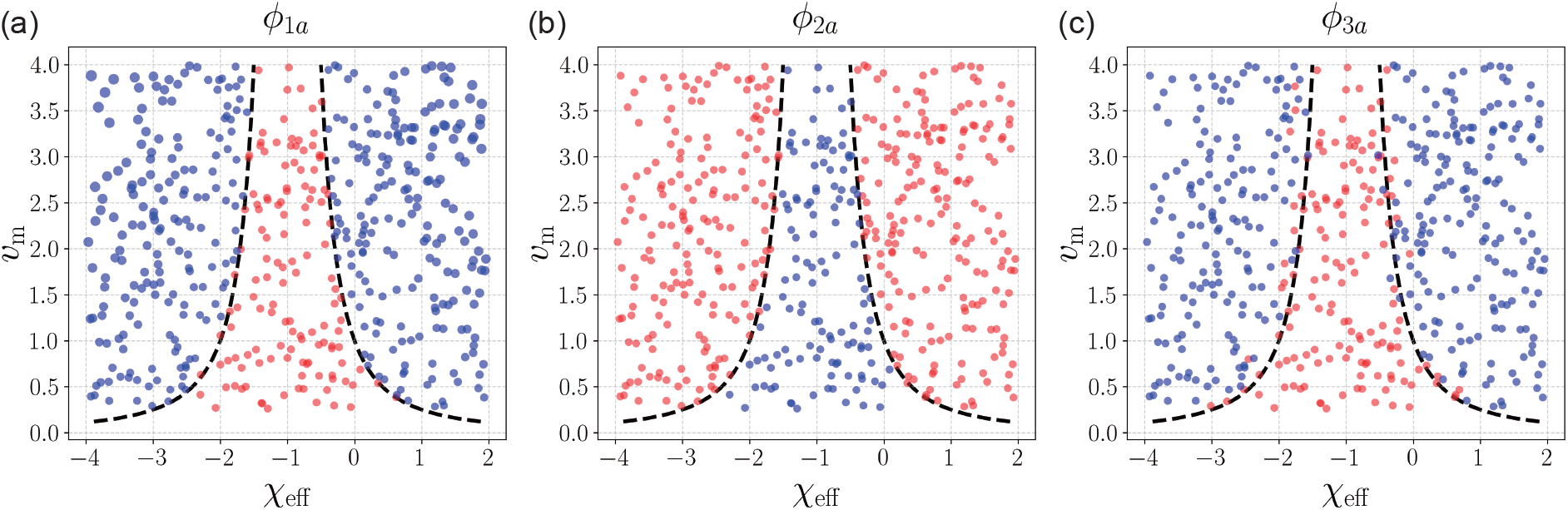
Linear responses of a multicomponent solution to macromolecule addition. (a) The sign of the change in the major-phase volume fraction of the 1st biomolecule (blue: negative; red: positive). Here, each point represents a type of macromolecule, with the x- and y-axis variables being its size *v*_*m*_ and effective interaction *χ*_eff_ with preexisting biomolecules, respectively. The black dashed lines are the predicted zero-level curves along which 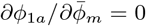. (b) The same as (a) but for the 2nd biomolecule. (c) The same as (a) but for the 3rd biomolecule. In this figure, the 1st and 3rd biomolecule species are enriched in the minor phase and the 2nd species is enriched in the major phase. For all the biomolecules, they share the same zero-level curves. In this figure, 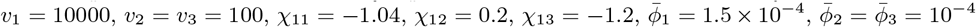, and the data are numerically computed from two states with 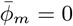 and 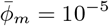.

The research was funded by the National Key Research and Development Program of China (2024YFA0919600), the National Natural Science Foundation of China (Grant No. 12474190) and Peking-Tsinghua Center for Life Sciences grants.

## Notes

### Competing Interest Statement

The authors have declared no competing interest.

## References

[1] Clifford P Brangwynne, Christian R Eckmann, David S Courson, Agata Rybarska, Carsten Hoege, Jöbin Gharakhani, Frank Jülicher, and Anthony A Hyman. Germline p granules are liquid droplets that localize by controlled dissolution/condensation. Science, 324(5935):1729–1732, 2009.

[2] Clifford P Brangwynne, Peter Tompa, and Rohit V Pappu. Polymer physics of intracellular phase transitions. Nature Physics, 11(11):899–904, 2015.

[3] Salman F Banani, Hyun O Lee, Anthony A Hyman, and Michael K Rosen. Biomolecular condensates: organizers of cellular biochemistry. Nature reviews Molecular cell biology, 18(5):285–298, 2017.

[4] Yongdae Shin and Clifford P Brangwynne. Liquid phase condensation in cell physiology and disease. Science, 357(6357):eaaf4382, 2017.

[5] Frank Jülicher and Christoph A Weber. Droplet physics and intracellular phase separation. Annual Review of Condensed Matter Physics, 15(1):237–261, 2024.

[6] Andrew S Lyon, William B Peeples, and Michael K Rosen. A framework for understanding the functions of biomolecular condensates across scales. Nature Reviews Molecular Cell Biology, 22(3):215–235, 2021.

[7] Lingyu Meng, Sheng Mao, and Jie Lin. Heterogeneous elasticity drives ripening and controls bursting kinetics of transcriptional condensates. Proceedings of the National Academy of Sciences, 121(12):e2316610121, 2024.

[8] Yichun Wu, Xing Wang, Lingyu Meng, Zhizhao Liao, Wei Ji, Peipei Zhang, Jie Lin, and Qiang Guo. Translation landscape of stress granules. Science Advances, 11(40):eady6859, 2025.

[9] Alain A. M. André and Evan Spruijt. Liquid–liquid phase separation in crowded environments. International Journal of Molecular Sciences, 21(16), 2020.

[10] Yuan Lin, David SW Protter, Michael K Rosen, and Roy Parker. Formation and maturation of phase-separated liquid droplets by rna-binding proteins. Molecular cell, 60(2):208–219, 2015.

[11] Salman F Banani, Allyson M Rice, William B Peeples, Yuan Lin, Saumya Jain, Roy Parker, and Michael K Rosen. Compositional control of phase-separated cellular bodies. Cell, 166(3):651–663, 2016.

[12] Shambaditya Saha, Christoph A. Weber, Marco Nousch, Omar Adame-Arana, Carsten Hoege, Marco Y. Hein, Erin Osborne-Nishimura, Julia Mahamid, Marcus Jahnel, Louise Jawerth, Andrej Pozniakovski, Christian R. Eckmann, Frank Jülicher, and Anthony A. Hyman. Polar positioning of phase-separated liquid compartments in cells regulated by an mrna competition mechanism. Cell, 166(6):1572–1584.e16, 2016.

[13] Jarrett Smith, Deepika Calidas, Helen Schmidt, Tu Lu, Dominique Rasoloson, and Geraldine Seydoux. Spatial patterning of p granules by rna-induced phase separation of the intrinsically-disordered protein meg-3. Elife, 5:e21337, 2016.

[14] Marina Feric, Nilesh Vaidya, Tyler S Harmon, Diana M Mitrea, Lian Zhu, Tiffany M Richardson, Richard W Kriwacki, Rohit V Pappu, and Clifford P Brangwynne. Coexisting liquid phases underlie nucleolar subcompartments. Cell, 165(7):1686–1697, 2016.

[15] Jordina Guillén-Boixet, Andrii Kopach, Alex S. Holehouse, Sina Wittmann, Marcus Jahnel, Raimund Schlüßler, Kyoohyun Kim, Irmela R.E.A. Trussina, Jie Wang, Daniel Mateju, Ina Poser, Shovamayee Maharana, Martine Ruer-Gruß, Doris Richter, Xiaojie Zhang, Young-Tae Chang, Jochen Guck, Alf Honigmann, Julia Mahamid, Anthony A. Hyman, Rohit V. Pappu, Simon Alberti, and Titus M. Franzmann. Rna-induced conformational switching and clustering of g3bp drive stress granule assembly by condensation. Cell, 181(2):346–361.e17, 2020.

[16] Amandine Molliex, Jamshid Temirov, Jihun Lee, Maura Coughlin, Anderson P Kanagaraj, Hong Joo Kim, Tanja Mittag, and J Paul Taylor. Phase separation by low complexity domains promotes stress granule assembly and drives pathological fibrillization. Cell, 163(1):123–133, 2015.

[17] Susanne Wegmann, Bahareh Eftekharzadeh, Katharina Tepper, Katarzyna M Zoltowska, Rachel E Bennett, Simon Dujardin, Pawel R Laskowski, Danny MacKenzie, Tarun Kamath, Caitlin Commins, et al. Tau protein liquid–liquid phase separation can initiate tau aggregation. The EMBO journal, 37(7):EMBJ201798049, 2018.

[18] Jie Wang, Jeong-Mo Choi, Alex S Holehouse, Hyun O Lee, Xiaojie Zhang, Marcus Jahnel, Shovamayee Maharana, Régis Lemaitre, Andrei Pozniakovsky, David Drechsel, et al. A molecular grammar governing the driving forces for phase separation of prion-like rna binding proteins. Cell, 174(3):688–699, 2018.

[19] Pilong Li, Sudeep Banjade, Hui-Chun Cheng, Soyeon Kim, Baoyu Chen, Liang Guo, Marc Llaguno, Javoris V Hollingsworth, David S King, Salman F Banani, et al. Phase transitions in the assembly of multivalent signalling proteins. Nature, 483(7389):336–340, 2012.

[20] Huaiying Zhang, Shana Elbaum-Garfinkle, Erin M. Langdon, Nicole Taylor, Patricia Occhipinti, Andrew A. Bridges, Clifford P. Brangwynne, and Amy S. Gladfelter. Rna controls polyq protein phase transitions. Molecular Cell, 60(2):220–230, 2015.

[21] Priya R. Banerjee, Anthony N. Milin, Mahdi Muhammad Moosa, Paulo L. Onuchic, and Ashok A. Deniz. Reentrant phase transition drives dynamic substructure formation in ribonucleoprotein droplets. Angewandte Chemie International Edition, 56(38):11354–11359, 2017.

[22] Shovamayee Maharana, Jie Wang, Dimitrios K. Papadopoulos, Doris Richter, Andrey Pozniakovsky, Ina Poser, Marc Bickle, Sandra Rizk, Jordina Guillén-Boixet, Titus M. Franzmann, Marcus Jahnel, Lara Marrone, Young-Tae Chang, Jared Sterneckert, Pavel Tomancak, Anthony A. Hyman, and Simon Alberti. Rna buffers the phase separation behavior of prion-like rna binding proteins. Science, 360(6391):918–921, 2018.

[23] Ibraheem Alshareedah, Taranpreet Kaur, Jason Ngo, Hannah Seppala, Liz-Audrey Djomnang Kounatse, Wei Wang, Mahdi Muhammad Moosa, and Priya R. Banerjee. Interplay between short-range attraction and longrange repulsion controls reentrant liquid condensation of ribonucleoprotein–rna complexes. Journal of the American Chemical Society, 141(37):14593–14602, 2019. PMID: 31437398.

[24] Paul J Flory. Thermodynamics of high polymer solutions. The Journal of chemical physics, 10(1):51–61, 1942.

[25] Maurice L Huggins. Some properties of solutions of long-chain compounds. The Journal of Physical Chemistry, 46(1):151–158, 1942.

[26] Joel Berry, Clifford P Brangwynne, and Mikko Haataja. Physical principles of intracellular organization via active and passive phase transitions. Reports on Progress in Physics, 81(4):046601, 2018.

[27] Michael Rubinstein and Ralph H Colby. Polymer physics. Oxford university press, 2003.

[28] Daoyuan Qian, Thomas C. T. Michaels, and Tuomas P. J. Knowles. Analytical solution to the Flory–Huggins model. The Journal of Physical Chemistry Letters, 13(33):7853–7860, 2022.

[29] Archishman Ghosh, Konstantinos Mazarakos, and Huan-Xiang Zhou. Three archetypical classes of macromolecular regulators of protein liquid–liquid phase separation. Proceedings of the National Academy of Sciences, 116(39):19474–19483, 2019.

[30] Ming-Tzo Wei, Shana Elbaum-Garfinkle, Alex S Holehouse, Carlos Chih-Hsiung Chen, Marina Feric, Craig B Arnold, Rodney D Priestley, Rohit V Pappu, and Clifford P Brangwynne. Phase behaviour of disordered proteins underlying low density and high permeability of liquid organelles. Nature chemistry, 9(11):1118–1125, 2017.

[31] Patrick M McCall, Kyoohyun Kim, Anna Shevchenko, Martine Ruer-Gruß, Jan Peychl, Jochen Guck, Andrej Shevchenko, Anthony A Hyman, and Jan Brugués. A label-free method for measuring the composition of multicomponent biomolecular condensates. Nature Chemistry, 17(12):1891–1902, 2025.

[32] Pierre-Gilles de Gennes. Scaling concepts in polymer physics. Cornell university press, 1979.

[33] Masao Doi and Sam F Edwards. The theory of polymer dynamics. Oxford University Press, 1988.

[34] Valery Nguemaha and Huan-Xiang Zhou. Liquid-liquid phase separation of patchy particles illuminates diverse effects of regulatory components on protein droplet formation. Scientific reports, 8(1):6728, 2018.

