## Supplemental Material for "Universal modulation of liquid-liquid phase separation by macromolecules"

Yunxiao Kan[1] and Jie Lin[2,3]
[1]Yuanpei College, Peking University, Beijing, China
[2]Center for Quantitative Biology, Peking University, Beijing, China
[3]Peking-Tsinghua Center for Life Sciences, Peking University, Beijing, China
(Dated: September 2, 2026)

#### A. FLORY-HUGGINS MODEL FOR THREE-COMPONENT SYSTEMS

We consider a three-component solution containing a scaffold protein with chain length $v_1$ (denoted as 1), a macromolecule (i.e., cosolute, denoted as 2), and a solvent (denoted as 3). We set the volume of the solvent, $v_0$, as the unit volume. The free energy density in unit of $k_BT/v_0$ can be expressed as follows:

$$f(\phi_1, \phi_2) = \frac{1}{v_1}\phi_1 \ln \phi_1 + \frac{1}{v_2}\phi_2 \ln \phi_2 + (1 - \phi_1 - \phi_2)\ln(1 - \phi_1 - \phi_2) + \frac{1}{2}\chi_{11}\phi_1^2 + \chi_{12}\phi_1\phi_2 + \frac{1}{2}\chi_{22}\phi_2^2, \tag{1}$$

where $v_1$ and $v_2$ are the volumes of the protein and the macromolecule, respectively. $\phi_1, \phi_2, \phi_3$ are the volume fractions of the three components, respectively, and satisfy $\phi_3 = 1 - \phi_1 - \phi_2$. $\chi_{11}, \chi_{12}, \chi_{22}$ represent the interaction parameters between protein-protein, protein-macromolecule, and macromolecule-macromolecule, respectively. Based on the free energy, we obtain the chemical potentials of the two species and the osmotic pressure of the solution:

$$\mu_1 = \frac{\partial f}{\partial \phi_1} = \frac{1}{v_1}\ln\phi_1 + \frac{1}{v_1} - \ln(1-\phi_1-\phi_2) - 1 + \chi_{11}\phi_1 + \chi_{12}\phi_2, \tag{2}$$

$$\mu_2 = \frac{\partial f}{\partial \phi_2} = \frac{1}{v_2}\ln\phi_2 + \frac{1}{v_2} - \ln(1-\phi_1-\phi_2) - 1 + \chi_{12}\phi_1 + \chi_{22}\phi_2, \tag{3}$$

$$\Pi = -f + \phi_1\mu_1 + \phi_2\mu_2 = \Big(\frac{1}{v_1} - 1\Big)\phi_1 + \Big(\frac{1}{v_2} - 1\Big)\phi_2 - \ln(1-\phi_1-\phi_2) + \frac{1}{2}\chi_{11}\phi_1^2 + \chi_{12}\phi_1\phi_2 + \frac{1}{2}\chi_{22}\phi_2^2. \tag{4}$$

We assume that in the absence of the macromolecule, the protein solution undergoes phase separation, forming a dense phase and a dilute phase. After adding the macromolecule, the solution still separates into two phases. We denote the dilute phase as 'a' and the dense phase as 'b'. At equilibrium, the volume fractions of each species in the two phases satisfy the following equations:

$$\begin{cases} \mu_{1a} - \mu_{1b} &= 0 & \text{(5a)} \\ \mu_{2a} - \mu_{2b} &= 0 & \text{(5b)} \\ \Pi_a - \Pi_b &= 0 & \text{(5c)} \\ f_a\phi_{1a} + (1-f_a)\phi_{1b} - \bar{\phi}_1 &= 0 & \text{(5d)} \\ f_a\phi_{2a} + (1-f_a)\phi_{2b} - \bar{\phi}_2 &= 0\ , & \text{(5e)} \end{cases}$$

where $f_a$ is the volume ratio of the dilute phase to the solution, and $\bar{\phi}_1, \bar{\phi}_2$ are the overall (i.e., average) volume fractions of the two solutes in the entire system. We also define $f_b \equiv 1 - f_a$ as the volume ratio of the dense phase. By solving this set of equations, we can determine $\phi_{1a}, \phi_{1b}, \phi_{2a}, \phi_{2b}$ and $f_a$, which can be considered as functions of the parameters $v_1, v_2, \chi_{11}, \chi_{12}, \chi_{22}, \bar{\phi}_1, \bar{\phi}_2$.

#### B. EFFECTS OF PARAMETER CHANGES ON PHASE SEPARATION

We study the effects of changes in model parameters on phase separation by calculating the derivatives of the protein volume fractions in the dilute and dense phases with respect to each parameter. In the following, we use the lower indexes 'a' and 'b' to represent variables associated with the dilute and dense phases, respectively. Let the parameter of interest be X, which can be any of $v_1, v_2, \chi_{11}, \chi_{12}, \chi_{22}, \bar{\phi}_1, \bar{\phi}_2$. Keeping other parameters constant, we perform a first-order expansion of Eq. (5) to obtain

$$\begin{pmatrix} \frac{\partial \mu_{1a}}{\partial \phi_{1a}} & -\frac{\partial \mu_{1b}}{\partial \phi_{1b}} & \frac{\partial \mu_{1a}}{\partial \phi_{2a}} & -\frac{\partial \mu_{1b}}{\partial \phi_{2b}} & 0 \\ \frac{\partial \mu_{2a}}{\partial \phi_{1a}} & -\frac{\partial \mu_{2b}}{\partial \phi_{1b}} & \frac{\partial \mu_{2a}}{\partial \phi_{2a}} & -\frac{\partial \mu_{2b}}{\partial \phi_{2b}} & 0 \\ \frac{\partial \Pi_{a}}{\partial \phi_{1a}} & -\frac{\partial \Pi_{b}}{\partial \phi_{1b}} & \frac{\partial \Pi_{a}}{\partial \phi_{2a}} & -\frac{\partial \Pi_{b}}{\partial \phi_{2b}} & 0 \\ f_a & 1-f_a & 0 & 0 & \phi_{1a}-\phi_{1b} \\ 0 & 0 & f_a & 1-f_a & \phi_{2a}-\phi_{2b} \end{pmatrix} \begin{pmatrix} \delta\phi_{1a} \\ \delta\phi_{1b} \\ \delta\phi_{2a} \\ \delta\phi_{2b} \\ \delta f_a \end{pmatrix} = \begin{pmatrix} -\frac{\partial \mu_{1a}}{\partial X} + \frac{\partial \mu_{1b}}{\partial X} \\ -\frac{\partial \mu_{2a}}{\partial X} + \frac{\partial \mu_{2b}}{\partial X} \\ -\frac{\partial \Pi_{a}}{\partial X} + \frac{\partial \Pi_{b}}{\partial X} \\ \frac{\partial \bar{\phi}_1}{\partial X} \\ \frac{\partial \bar{\phi}_2}{\partial X} \end{pmatrix} \delta X. \tag{6}$$

Substituting

$$\frac{\partial \Pi}{\partial \phi_i} = \frac{\partial(-f+\phi_1\mu_1+\phi_2\mu_2)}{\partial \phi_i} = \phi_1\frac{\partial \mu_1}{\partial \phi_i} + \phi_2\frac{\partial \mu_2}{\partial \phi_i}, \quad \frac{\partial \mu_i}{\partial \phi_j} = \frac{\partial^2 f}{\partial \phi_i \partial \phi_j} \equiv f_{ij}, \tag{7}$$

into (6), we get

$$\begin{pmatrix} f_{11}^a & -f_{11}^b & f_{12}^a & -f_{12}^b & 0 \\ f_{12}^a & -f_{12}^b & f_{22}^a & -f_{22}^b & 0 \\ \phi_{1a}f_{11}^a+\phi_{2a}f_{12}^a & -(\phi_{1b}f_{11}^b+\phi_{2b}f_{12}^b) & \phi_{1a}f_{12}^a+\phi_{2a}f_{22}^a & -(\phi_{1b}f_{12}^b+\phi_{2b}f_{22}^b) & 0 \\ f_a & 1-f_a & 0 & 0 & \phi_{1a}-\phi_{1b} \\ 0 & 0 & f_a & 1-f_a & \phi_{2a}-\phi_{2b} \end{pmatrix} \begin{pmatrix} \delta\phi_{1a} \\ \delta\phi_{1b} \\ \delta\phi_{2a} \\ \delta\phi_{2b} \\ \delta f_a \end{pmatrix} = \vec{\beta}_X \delta X, \tag{8}$$

where $\vec{\beta}_X$ is the vector of expansion coefficients with respect to X on the right side of Eq. (6). Specifically,

$$\vec{\beta}_{v_2} = -\frac{1}{v_2^2}\begin{pmatrix} 0 \\ \ln\frac{\phi_{2b}}{\phi_{2a}} \\ \phi_{2b}-\phi_{2a} \\ 0 \\ 0 \end{pmatrix}, \vec{\beta}_{\chi_{11}} = \begin{pmatrix} \phi_{1b}-\phi_{1a} \\ 0 \\ \frac{1}{2}(\phi_{1b}^2-\phi_{1a}^2) \\ 0 \\ 0 \end{pmatrix}, \vec{\beta}_{\chi_{12}} = \begin{pmatrix} \phi_{2b}-\phi_{2a} \\ \phi_{1b}-\phi_{1a} \\ \phi_{1b}\phi_{2b}-\phi_{1a}\phi_{2a} \\ 0 \\ 0 \end{pmatrix}, \vec{\beta}_{\chi_{22}} = \begin{pmatrix} 0 \\ \phi_{2b}-\phi_{2a} \\ \frac{1}{2}(\phi_{2b}^2-\phi_{2a}^2) \\ 0 \\ 0 \end{pmatrix}, \vec{\beta}_{\bar{\phi}_2} = \begin{pmatrix} 0 \\ 0 \\ 0 \\ 0 \\ 1 \end{pmatrix}. \tag{9}$$

Let $\mathbf{M}$ be the matrix on the left side of Eq. (8), which is invertible as we show later; formally, we get

$$\begin{pmatrix} \frac{\partial \phi_{1a}}{\partial X} \\ \frac{\partial \phi_{1b}}{\partial X} \\ \frac{\partial \phi_{2a}}{\partial X} \\ \frac{\partial \phi_{2b}}{\partial X} \\ \frac{\partial f_a}{\partial X} \end{pmatrix} = \mathbf{M}^{-1}\vec{\beta}_X. \tag{10}$$

Solving for the inverse of $\mathbf{M}$ is difficult. Nevertheless, we can use Cramer's rule to bypass this problem. Specifically, if the quantity to be solved is in the i-th row on the left side of Eq. (10), we replace the i-th column of matrix $\mathbf{M}$ with $\vec{\beta}_X$ to get matrix $\mathbf{M}'$, and then find the ratio of the determinant of $\mathbf{M}'$ to the determinant of $\mathbf{M}$. For example,

$$\frac{\partial\phi_{1a}}{\partial\bar{\phi}_2}=\frac{|\mathbf{M}'|_{\bar{\phi}_2}}{|\mathbf{M}|}=\begin{vmatrix}0 & -f^b_{11} & f^a_{12} & -f^b_{12} & 0\\ 0 & -f^b_{12} & f^a_{22} & -f^b_{22} & 0\\ 0 & -(\phi_{1b}f^b_{11}+\phi_{2b}f^b_{12}) & \phi_{1a}f^a_{12}+\phi_{2a}f^a_{22} & -(\phi_{1b}f^b_{12}+\phi_{2b}f^b_{22}) & 0\\ 0 & 1-f_a & 0 & 0 & \phi_{1a}-\phi_{1b}\\ 1 & 0 & f_a & 1-f_a & \phi_{2a}-\phi_{2b}\end{vmatrix}\cdot|\mathbf{M}|^{-1}. \tag{11}$$

We now prove that the determinant of matrix $\mathbf{M}$ is positive. Direct calculation yields

$$\begin{aligned}|\mathbf{M}|=&(1-f_a)\cdot\begin{pmatrix}\phi_{1a}-\phi_{1b} & \phi_{2a}-\phi_{2b}\end{pmatrix}\begin{pmatrix}f^b_{11} & f^b_{12}\\ f^b_{12} & f^b_{22}\end{pmatrix}\begin{pmatrix}\phi_{1a}-\phi_{1b}\\ \phi_{2a}-\phi_{2b}\end{pmatrix}\cdot\begin{vmatrix}f^a_{11} & f^a_{12}\\ f^a_{12} & f^a_{22}\end{vmatrix}\\ &+f_a\cdot\begin{pmatrix}\phi_{1a}-\phi_{1b} & \phi_{2a}-\phi_{2b}\end{pmatrix}\begin{pmatrix}f^a_{11} & f^a_{12}\\ f^a_{12} & f^a_{22}\end{pmatrix}\begin{pmatrix}\phi_{1a}-\phi_{1b}\\ \phi_{2a}-\phi_{2b}\end{pmatrix}\cdot\begin{vmatrix}f^b_{11} & f^b_{12}\\ f^b_{12} & f^b_{22}\end{vmatrix}.\end{aligned} \tag{12}$$

We remark a trick in obtaining the above equation. One can find $|\mathbf{M}|$ with $f_a=0$ and $f_a=1$ first, using Laplacian theorem. Since $|\mathbf{M}|$ must be a linear function of $f_a$, the determinant with $0<f_a<1$ must be a linear interpolation of the determinants with $f_a=0$ and $f_a=1$.

We note that $|\mathbf{M}|$ has two terms proportional to $f_a$ and $1-f_a$, respectively. Both of them are quadratic forms of the Hessian matrix multiplied by the determinant of the other phase's Hessian matrix. In equilibrium, both phases are locally stable, and their Hessian matrices are positive definite, so the quadratic forms and determinants of the Hessian matrices are also positive.

#### C. SOME GENERAL CONCLUSIONS ON THE EFFECTS OF PARAMETER CHANGES ON PHASE SEPARATION

We next study whether a parameter change promotes or suppresses phase separation. We use $\partial\phi_{1a}/\partial X$ to determine the effect of parameter $X$ on the protein concentration in the dilute phase. A negative value indicates promotion of phase separation, while a positive value indicates suppression of phase separation. Since the denominator is positive, if we want to find the sign of $\partial\phi_{1a}/\partial X$, we only need to consider the numerator, $|\mathbf{M}'|$. In the following, we analyze the first four variables in Eq. (9) in three scenarios. Later, we analyze the effect of $\bar{\phi}_2$ separately.

##### C1. Case where $v_2=1, \chi_{12}=\chi_{22}=0$

In this case, the macromolecule and solvent are identical. Using this state as the reference state, we tune the properties of the macromolecule and discuss how the solution's phase separation changes. Equilibrium of the chemical potential of the macromolecule [Eq. (5b)] in this scenario becomes

$$\ln\phi_{2a}-\ln\phi_{3a}=\ln\phi_{2b}-\ln\phi_{3b}\Rightarrow\frac{\phi_{3a}}{\phi_{2a}}=\frac{\phi_{3b}}{\phi_{2b}}=\frac{\bar{\phi}_3}{\bar{\phi}_2}\equiv\lambda-1. \tag{13}$$

Here, we introduce $\lambda$ for convenience. We also have

$$f_{12}=\frac{1}{\phi_3}+\chi_{12}=\frac{1}{\phi_3}, \tag{14}$$

$$f_{22}=\frac{1}{v_2\phi_2}+\frac{1}{\phi_3}+\chi_{22}=\frac{\lambda}{\phi_3}, \tag{15}$$

$$\phi_1 f_{12}+\phi_2 f_{22}=\frac{1}{\phi_3}. \tag{16}$$

First, we consider $\partial\phi_{1a}/\partial v_2$:

$$|\mathbf{M}'|_{v_2} = \begin{vmatrix} 0 & -f^b_{11} & \dfrac{1}{\phi_{3a}} & -\dfrac{1}{\phi_{3b}} & 0 \\ -\frac{1}{v_2^2}\ln\left(\dfrac{\phi_{2b}}{\phi_{2a}}\right) & -\dfrac{1}{\phi_{3b}} & \dfrac{\lambda}{\phi_{3a}} & -\dfrac{\lambda}{\phi_{3b}} & 0 \\ -\frac{1}{v_2^2}(\phi_{2b}-\phi_{2a}) & -\phi_{1b}f^b_{11} - \dfrac{1}{\lambda-1} & \dfrac{1}{\phi_{3a}} & -\dfrac{1}{\phi_{3b}} & 0 \\ 0 & 1-f_a & 0 & 0 & \phi_{1a}-\phi_{1b} \\ 0 & 0 & f_a & 1-f_a & \phi_{2a}-\phi_{2b} \end{vmatrix} \tag{17}$$
$$= -\frac{1}{v_2^2}(\phi_{1b}-\phi_{1a})\Big(\frac{1-f_a}{\phi_{3a}} + \frac{f_a}{\phi_{3b}}\Big)\Big(\phi_{2a} - \phi_{2b} - \phi_{2b}\ln\frac{\phi_{2a}}{\phi_{2b}}\Big)\Big(\lambda f^b_{11} - \frac{1}{\phi_{3b}}\Big).$$

Obviously, $\phi_{1b} - \phi_{1a}$ and $(1-f_a)/\phi_{3a} + f_a/\phi_{3b}$ are greater than zero:

$$\phi_{2a} - \phi_{2b} - \phi_{2b}\ln\frac{\phi_{2a}}{\phi_{2b}} > \phi_{2a} - \phi_{2b} - \phi_{2b}\Big(\frac{\phi_{2a}}{\phi_{2b}} - 1\Big) = 0. \tag{18}$$

The determinant of the Hessian matrix is also greater than zero:

$$\begin{pmatrix} f^b_{11} & f^b_{12} \\ f^b_{12} & f^b_{22} \end{pmatrix} = f^b_{11}\frac{\lambda}{\phi_{3b}} - \frac{1}{\phi_{3b}^2} = \frac{1}{\phi_{3b}}\Big(\lambda f^b_{11} - \frac{1}{\phi_{3b}}\Big) > 0 \Rightarrow \lambda f^b_{11} - \frac{1}{\phi_{3b}} > 0. \tag{19}$$

Therefore, in this case, $\partial\phi_{1a}/\partial v_2 < 0$. This means that for a macromolecule that is identical to the solvent in terms of interaction, if its volume is larger than the solvent ($v_2 > 1$), it promotes phase separation; if its volume is smaller than the solvent ($v_2 < 1$), it suppresses phase separation.

Next, we consider $\partial\phi_{1a}/\partial\chi_{11}$:

$$|\mathbf{M}'|_{\chi_{11}} = \begin{vmatrix} \phi_{1b}-\phi_{1a} & -f^b_{11} & \dfrac{1}{\phi_{3a}} & -\dfrac{1}{\phi_{3b}} & 0 \\ 0 & -\dfrac{1}{\phi_{3b}} & \dfrac{\lambda}{\phi_{3a}} & -\dfrac{\lambda}{\phi_{3b}} & 0 \\ \frac{1}{2}(\phi_{1b}^2-\phi_{1a}^2) & -\phi_{1b}f^b_{11} - \dfrac{1}{\lambda-1} & \dfrac{1}{\phi_{3a}} & -\dfrac{1}{\phi_{3b}} & 0 \\ 0 & 1-f_a & 0 & 0 & \phi_{1a}-\phi_{1b} \\ 0 & 0 & f_a & 1-f_a & \phi_{2a}-\phi_{2b} \end{vmatrix} \tag{20}$$
$$= \frac{1}{2}(\phi_{1b}-\phi_{1a})^3\Big(\frac{1-f_a}{\phi_{3a}} + \frac{f_a}{\phi_{3b}}\Big)\Big(\lambda f^b_{11} - \frac{1}{\phi_{3b}}\Big) > 0.$$

The conclusion is obvious: stronger attractive interaction between protein molecules (i.e., smaller $\chi_{11}$) promotes phase separation.

Then, we consider $\partial\phi_{1a}/\partial\chi_{12}$. In this case, $\phi_2 = (1-\phi_1)/\lambda$, so $\phi_{1a} < \phi_{1b} \Rightarrow \phi_{2a} > \phi_{2b}$. Therefore,

$$|\mathbf{M}'|_{\chi_{12}} = \begin{vmatrix} \phi_{2b}-\phi_{2a} & -f^b_{11} & \dfrac{1}{\phi_{3a}} & -\dfrac{1}{\phi_{3b}} & 0 \\ \phi_{1b}-\phi_{1a} & -\dfrac{1}{\phi_{3b}} & \dfrac{\lambda}{\phi_{3a}} & -\dfrac{\lambda}{\phi_{3b}} & 0 \\ \phi_{1b}\phi_{2b}-\phi_{1a}\phi_{2a} & -\phi_{1b}f^b_{11} - \dfrac{1}{\lambda-1} & \dfrac{1}{\phi_{3a}} & -\dfrac{1}{\phi_{3b}} & 0 \\ 0 & 1-f_a & 0 & 0 & \phi_{1a}-\phi_{1b} \\ 0 & 0 & f_a & 1-f_a & \phi_{2a}-\phi_{2b} \end{vmatrix} \tag{21}$$
$$= (\phi_{1b}-\phi_{1a})^2(\phi_{2b}-\phi_{2a})\Big(\frac{1-f_a}{\phi_{3a}} + \frac{f_a}{\phi_{3b}}\Big)\Big(\lambda f^b_{11} - \frac{1}{\phi_{3b}}\Big) < 0.$$

This can be understood as: when $\chi_{12}$ increases, the repulsion between protein and macromolecule strengthens. Since there are more macromolecules in the dilute phase than in the dense phase, protein molecules are repelled by macromolecules in the dilute phase into the dense phase, thereby promoting phase separation.

Finally, we consider $\partial\phi_{1a}/\partial\chi_{22}$:

$$|\mathbf{M}'|_{\chi_{22}} = \begin{vmatrix} 0 & -f_{11}^b & \frac{1}{\phi_{3a}} & -\frac{1}{\phi_{3b}} & 0 \\ \phi_{2b}-\phi_{2a} & -\frac{1}{\phi_{3b}} & \frac{\lambda}{\phi_{3a}} & -\frac{\lambda}{\phi_{3b}} & 0 \\ \frac{1}{2}(\phi_{2b}^2-\phi_{2a}^2) & -\phi_{1b}f_{11}^b - \frac{1}{\lambda-1} & \frac{1}{\phi_{3a}} & -\frac{1}{\phi_{3b}} & 0 \\ 0 & 1-f_a & 0 & 0 & \phi_{1a}-\phi_{1b} \\ 0 & 0 & f_a & 1-f_a & \phi_{2a}-\phi_{2b} \end{vmatrix} = \frac{1}{2}(\phi_{1b}-\phi_{1a})(\phi_{2b}-\phi_{2a})^2\Big(\frac{1-f_a}{\phi_{3a}}+\frac{f_a}{\phi_{3b}}\Big)\Big(\lambda f_{11}^b - \frac{1}{\phi_{3b}}\Big) > 0. \quad (22)$$

##### C2. Case where $f_a = 1$

In this case, the system is right at the boundary in the parameter space between the single-phase region and the two-phase region, i.e., at the edge of forming condensates. We substitute $f_a = 1$ into the expression for $|\mathbf{M}'|$ to see its sign.

First, we consider $\partial\phi_{1a}/\partial v_2$, which is negative, showing that large volume macromolecule promotes phase separation:

$$|\mathbf{M}'|_{v_2} = \begin{vmatrix} 0 & -f_{11}^b & f_{12}^a & -f_{12}^b & 0 \\ -\frac{1}{v_2^2}\ln\Big(\frac{\phi_{2b}}{\phi_{2a}}\Big) & -f_{12}^b & f_{22}^a & -f_{22}^b & 0 \\ -\frac{1}{v_2^2}(\phi_{2b}-\phi_{2a}) & -(\phi_{1b}f_{11}^b+\phi_{2b}f_{12}^b) & \phi_{1a}f_{12}^a+\phi_{2a}f_{22}^a & -(\phi_{1b}f_{12}^b+\phi_{2b}f_{22}^b) & 0 \\ 0 & 0 & 0 & 0 & \phi_{1a}-\phi_{1b} \\ 0 & 0 & 1 & 0 & \phi_{2a}-\phi_{2b} \end{vmatrix} = -\frac{1}{v_2^2}(\phi_{1b}-\phi_{1a})\Big(\phi_{2a}-\phi_{2b}-\phi_{2b}\ln\frac{\phi_{2a}}{\phi_{2b}}\Big)\begin{vmatrix} f_{11}^b & f_{12}^b \\ f_{12}^b & f_{22}^b \end{vmatrix} < 0. \quad (23)$$

Next, we consider $\partial\phi_{1a}/\partial\chi_{11}$:

$$|\mathbf{M}'|_{\chi_{11}} = \frac{1}{2}(\phi_{1b}-\phi_{1a})^3\begin{vmatrix} f_{11}^b & f_{12}^b \\ f_{12}^b & f_{22}^b \end{vmatrix} > 0, \quad (24)$$

which is positive, showing that attraction between protein molecules promotes phase separation, while repulsion suppresses phase separation.

Then, we consider $\partial\phi_{1a}/\partial\chi_{12}$:

$$|\mathbf{M}'|_{\chi_{12}} = (\phi_{1b}-\phi_{1a})^2(\phi_{2b}-\phi_{2a})\begin{vmatrix} f_{11}^b & f_{12}^b \\ f_{12}^b & f_{22}^b \end{vmatrix}. \quad (25)$$

The sign of the result depends on $\phi_{2a}$ and $\phi_{2b}$. When $\phi_{2a} > \phi_{2b}$, macromolecule is enriched in the dilute phase, so stronger repulsion between macromolecule and protein enriches protein in the dense phase. Conversely, when $\phi_{2a} < \phi_{2b}$, macromolecule is enriched in the dense phase, so stronger attraction between macromolecule and protein enriches protein in the dense phase.

Finally, we consider $\partial\phi_{1a}/\partial\chi_{22}$:

$$|\mathbf{M}'|_{\chi_{22}} = \frac{1}{2}(\phi_{1b}-\phi_{1a})(\phi_{2b}-\phi_{2a})^2 \begin{vmatrix} f_{11}^b & f_{12}^b \\ f_{12}^b & f_{22}^b \end{vmatrix} > 0. \tag{26}$$

Interestingly, this implies that making the macromolecule-macromolecule interaction more repulsive suppresses phase separation.

##### C3. Effects of $\bar{\phi}_2$ on phase separation

The derivatives to $\bar{\phi}_2$ has a clear physical meaning: what happens to the phase separation of the solution when a small amount of macromolecule is added?

$$\begin{aligned} |\mathbf{M}'|_{\bar{\phi}_2} &= \begin{vmatrix} 0 & -f_{11}^b & f_{12}^a & -f_{12}^b & 0 \\ 0 & -f_{12}^b & f_{22}^a & -f_{22}^b & 0 \\ 0 & -(\phi_{1b}f_{11}^b+\phi_{2b}f_{12}^b) & \phi_{1a}f_{12}^a+\phi_{2a}f_{22}^a & -(\phi_{1b}f_{12}^b+\phi_{2b}f_{22}^b) & 0 \\ 0 & 1-f_a & 0 & 0 & \phi_{1a}-\phi_{1b} \\ 1 & 0 & f_a & 1-f_a & \phi_{2a}-\phi_{2b} \end{vmatrix} \\ &= (\phi_{1b}-\phi_{1a})((\phi_{1a}-\phi_{1b})f_{12}^a+(\phi_{2a}-\phi_{2b})f_{22}^a) \begin{vmatrix} f_{11}^b & f_{12}^b \\ f_{12}^b & f_{22}^b \end{vmatrix}. \end{aligned} \tag{27}$$

Interestingly, the expression for $|\mathbf{M}'|$ does not contain $f_a$, so the sign of $\partial\phi_{1a}/\partial\bar{\phi}_2$ is not affected by $f_a$ (however, its value is still affected by $f_a$, because the denominator $|\mathbf{M}|$ still contains $f_a$). The sign of $\partial\phi_{1a}/\partial\bar{\phi}_2$ depends on the following expression:

$$(\phi_{1a}-\phi_{1b})f_{12}^a+(\phi_{2a}-\phi_{2b})f_{22}^a = (\phi_{1a}-\phi_{1b})\Big(\frac{1}{\phi_{3a}}+\chi_{12}\Big)+(\phi_{2a}-\phi_{2b})\Big(\frac{1}{v_2\phi_{2a}}+\frac{1}{\phi_{3a}}+\chi_{22}\Big). \tag{28}$$

We consider a simpler case: $\chi_{12}=\chi_{22}=0$. Eq. (5b) becomes

$$\frac{1}{v_2}\ln\phi_{2a}-\ln\phi_{3a} = \frac{1}{v_2}\ln\phi_{2b}-\ln\phi_{3b} \Rightarrow \Big(\frac{\phi_{3b}}{\phi_{3a}}\Big)^{v_2} = \frac{\phi_{2b}}{\phi_{2a}}. \tag{29}$$

Let $t = 1-\phi_{3b}/\phi_{3a} > 0$, we have

$$(\phi_{1a}-\phi_{1b})f_{12}^a+(\phi_{2a}-\phi_{2b})f_{22}^a = \frac{\phi_{3b}-\phi_{3a}}{\phi_{3a}}+\frac{\phi_{2a}-\phi_{2b}}{v_2\phi_{2a}} = \frac{1}{v_2}(1-v_2t-(1-t)^{v_2}). \tag{30}$$

When $v_2>1$, $(1-t)^{v_2} > 1-v_2t$, $\partial\phi_{1a}/\partial\bar{\phi}_2 < 0$, phase separation is promoted; when $v_2=1$, $(1-t)^{v_2}=1-tv_2$, $\partial\phi_{1a}/\partial\bar{\phi}_2 = 0$, phase separation is not affected; when $v_2<1$, $(1-t)^{v_2} < 1-tv_2$, $\partial\phi_{1a}/\partial\bar{\phi}_2 > 0$, phase separation is suppressed. This conclusion is consistent with our previous discussion of $\partial\phi_{1a}/\partial v_2$, Eq. (17).

##### C4. Relationship between $\partial\phi_s/\partial\bar{\phi}_2$ and $(\partial\phi_{1a}/\partial\bar{\phi}_2)|_{f_a=1}$

We introduce $\phi_s$ as the saturation volume fraction of the protein, i.e., the minimum $\bar{\phi}_1$ to induce condensate formation starting from a homogeneous protein solution. By definition, we have $\phi_s = \phi_{1a}|_{f_a=1}$. However, we remark that $\partial\phi_s/\partial\bar{\phi}_2$ and $(\partial\phi_{1a}/\partial\bar{\phi}_2)|_{f_a=1}$ are generally different as we explain in the following.

$\partial\phi_s/\partial\bar{\phi}_2$ is defined as the change of saturation concentration with $\bar{\phi}_2$ so that $f_a$ is kept constant ($f_a=1$) when taking the derivative. Meanwhile, $\partial\phi_{1a}/\partial\bar{\phi}_2$ is defined as the change of dilute phase concentration with $\bar{\phi}_2$, and the average volume fraction of the protein, $\bar{\phi}_1$, is kept constant when taking the derivative. We derive the expression for $\partial\phi_s/\partial\bar{\phi}_2$ as follows. Starting from Eq. (5), we ignore equation (5d) because $\bar{\phi}_1$ is changing now. We perform a first-order expansion, this time keeping $f_a$ constant ($f_a = 1$):

$$\begin{pmatrix} f^a_{11} & -f^b_{11} & f^a_{12} & -f^b_{12} \\ f^a_{12} & -f^b_{12} & f^a_{22} & -f^b_{22} \\ \phi_{1a}f^a_{11} + \phi_{2a}f^a_{12} & -(\phi_{1b}f^b_{11} + \phi_{2b}f^b_{12}) & \phi_{1a}f^a_{12} + \phi_{2a}f^a_{22} & -(\phi_{1b}f^b_{12} + \phi_{2b}f^b_{22}) \\ 0 & 0 & 1 & 0 \end{pmatrix} \begin{pmatrix} \delta\phi_s \\ \delta\phi_{1b} \\ \delta\phi_{2a} \\ \delta\phi_{2b} \end{pmatrix} = \begin{pmatrix} 0 \\ 0 \\ 0 \\ \delta\bar{\phi}_2 \end{pmatrix}. \tag{31}$$

Solve this equation and we get

$$\frac{\partial\phi_s}{\partial\bar{\phi}_2} = -\frac{(\phi_{1a} - \phi_{1b})f^a_{12} + (\phi_{2a} - \phi_{2b})f^a_{22}}{(\phi_{1a} - \phi_{1b})f^a_{11} + (\phi_{2a} - \phi_{2b})f^a_{12}}. \tag{32}$$

Comparing with

$$\left.\frac{\partial\phi_{1a}}{\partial\bar{\phi}_2}\right|_{f_a=1} = \frac{(\phi_{1b} - \phi_{1a})((\phi_{1a} - \phi_{1b})f^a_{12} + (\phi_{2a} - \phi_{2b})f^a_{22})}{\begin{pmatrix} \phi_{1a} - \phi_{1b} & \phi_{2a} - \phi_{2b} \end{pmatrix} \begin{pmatrix} f^a_{11} & f^a_{12} \\ f^a_{12} & f^a_{22} \end{pmatrix} \begin{pmatrix} \phi_{1a} - \phi_{1b} \\ \phi_{2a} - \phi_{2b} \end{pmatrix}}, \tag{33}$$

we have

$$\frac{1}{\left.\dfrac{\partial\phi_{1a}}{\partial\bar{\phi}_2}\right|_{f_a=1}} = \frac{1}{\dfrac{\partial\phi_s}{\partial\bar{\phi}_2}} - \frac{\phi_{2b} - \phi_{2a}}{\phi_{1b} - \phi_{1a}}. \tag{34}$$

The last term on the right side of Eq. (34) is simply the slope of the tie line. Interestingly, if $\bar{\phi}_2 = 0$, $(\partial\phi_{1a}/\partial\bar{\phi}_2)|_{f_a=1} = \partial\phi_s/\partial\bar{\phi}_2$, which has the geometrical meaning as we show in the inset of Fig. 2b in the main text.

#### D. BEHAVIORS IN THE LIMIT $\bar{\phi}_2 \to 0$

##### D1. Partition of macromolecule in the two phases in the limit $\bar{\phi}_2 \to 0$

In the limit $\bar{\phi}_2 \to 0$, the system only contains protein and solvent. Therefore, we can use this binary system as a reference state and expand the equations for small $\bar{\phi}_2$. The free energy of a binary system is

$$f(\phi_1) = \frac{1}{v_1}\phi_1 \ln\phi_1 + (1 - \phi_1)\ln(1 - \phi_1) + \frac{1}{2}\chi_{11}\phi_1^2. \tag{35}$$

At equilibrium, the conditions on the chemical potential [Eq. (5a)], osmotic pressure [Eq. (5c)], and material conservation [Eq. (5d)] are satisfied in the limit $\bar{\phi}_2 \to 0$ as

$$\begin{cases} \dfrac{1}{v_1}\ln\phi_{1a} - \ln(1 - \phi_{1a}) + \chi_{11}\phi_{1a} = \dfrac{1}{v_1}\ln\phi_{1b} - \ln(1 - \phi_{1b}) + \chi_{11}\phi_{1b} & \text{(36a)} \\ \left(\dfrac{1}{v_1} - 1\right)\phi_{1a} - \ln(1 - \phi_{1a}) + \dfrac{1}{2}\chi_{11}\phi_{1a}^2 = \left(\dfrac{1}{v_1} - 1\right)\phi_{1b} - \ln(1 - \phi_{1b}) + \dfrac{1}{2}\chi_{11}\phi_{1b}^2. & \text{(36b)} \end{cases}$$

Based on this, $\phi_{1a}, \phi_{3a} = 1 - \phi_{1a}, \phi_{1b}, \phi_{3b} = 1 - \phi_{1b}$ can be determined. At the same time, $f_a$ can also be determined by mass conservation:

$$f_a\phi_{1a} + (1 - f_a)\phi_{1b} = \bar{\phi}_1. \tag{37}$$

After finding the reference state, we consider the volume fractions of the macromolecule in the two phases in the limit $\bar{\phi}_2 \to 0$, that is, adding an infinitesimal amount of the macromolecule to the system. Given $\phi_{2a}, \phi_{2b}$ and $\bar{\phi}_2$ as infinitesimals of the same order, we solve Eq. (5b) and Eq. (5e), neglecting higher-order infinitesimals:

$$\ln\frac{\phi_{2b}}{\phi_{2a}} = -v_2\Big(\chi_{12}(\phi_{1b}-\phi_{1a}) - \ln\frac{\phi_{3b}}{\phi_{3a}}\Big), \tag{38}$$

$$f_a\frac{\phi_{2a}}{\bar{\phi}_2} + (1-f_a)\frac{\phi_{2b}}{\bar{\phi}_2} = 1, \tag{39}$$

from which we find $\phi_{2a}/\bar{\phi}_2$ and $\phi_{2b}/\bar{\phi}_2$.

##### D2. Signs of derivatives in the limit $\bar{\phi}_2 \to 0$

We first compute $|\mathbf{M}|$ in the limit $\bar{\phi}_2 \to 0$ and only keep the most leading term,

$$|\mathbf{M}| \approx (\phi_{1b}-\phi_{1a})^2 f_{11}^a f_{11}^b \frac{\bar{\phi}_2}{v_2\phi_{2a}\phi_{2b}}, \tag{40}$$

which scales as $\bar{\phi}_2^{-1}$, where

$$f_{11} = \frac{1}{v_1\phi_1} + \frac{1}{\phi_3} + \chi_{11} > 0. \tag{41}$$

First, we consider $v_2$. When $f_a = 0$,

$$|\mathbf{M}'|_{v_2} = -\frac{1}{v_2^2}(\phi_{1b}-\phi_{1a})f_{11}^b\Big((\phi_{1b}-\phi_{1a})\Big(\frac{1}{\phi_{3a}}+\chi_{12}\Big)\ln\Big(\frac{\phi_{2a}}{\phi_{2b}}\Big) + \frac{1}{v_2}\Big(1+\ln\frac{\phi_{2b}}{\phi_{2a}} - \frac{\phi_{2b}}{\phi_{2a}}\Big)\Big). \tag{42}$$

This expression is of order unity. When $f_a = 1$,

$$|\mathbf{M}'|_{v_2} = -\frac{1}{v_2^3}(\phi_{1b}-\phi_{1a})f_{11}^b\Big(\frac{\phi_{2a}}{\phi_{2b}} - 1 - \ln\frac{\phi_{2a}}{\phi_{2b}}\Big) < 0, \tag{43}$$

which is also of order unity. We note that since $|\mathbf{M}'|_{v_2}$ is a linear function of $f_a$ [Eqs. (8, 9)], it must be of order unity for any $f_a$. Subsequently, after dividing by $|\mathbf{M}|$, which scale as $\bar{\phi}_2^{-1}$, $(\partial\phi_{1a}/\partial v_2)|_{\bar{\phi}_2\to 0}$ must be a first-order infinitesimal for any $f_a$.

Next, we consider $\chi_{11}$:

$$|\mathbf{M}'|_{\chi_{11}} = \frac{1}{2}(\phi_{1b}-\phi_{1a})^3 f_{11}^b \frac{\bar{\phi}_2}{v_2\phi_{2a}\phi_{2b}}, \tag{44}$$

so we have

$$\frac{\partial\phi_{1a}}{\partial\chi_{11}}\bigg|_{\bar{\phi}_2\to 0} = \frac{|\mathbf{M}'|_{\chi_{11}}}{|\mathbf{M}|} = \frac{\phi_{1b}-\phi_{1a}}{2f_{11}^a}, \tag{45}$$

which is obviously independent of $\bar{\phi}_2$ and the properties of the macromolecule.

Then we consider $\chi_{12}$:

$$f_a = 0, |\mathbf{M}'|_{\chi_{12}} = -(\phi_{1b}-\phi_{1a})^3 f_{11}^b\Big(\chi_{12}+\frac{1}{\phi_{3a}}\Big), \tag{46}$$

$$f_a = 1, |\mathbf{M}'|_{\chi_{12}} = \frac{1}{v_2}(\phi_{1b}-\phi_{1a})^2 f_{11}^b\Big(1-\frac{\phi_{2a}}{\phi_{2b}}\Big). \tag{47}$$

These expressions are of order unity. Since $|\mathbf{M}'|_{\chi_{12}}$ is a linear function of $f_a$, it must be of order unity for any $f_a$, so $(\partial\phi_{1a}/\partial\chi_{12})|_{\bar{\phi}_2\to 0}$ must be a first-order infinitesimal for any $f_a$.

Finally, we consider $\chi_{22}$:

$$f_a = 0, |\mathbf{M}'|_{\chi_{22}} = -(\phi_{1b}-\phi_{2b})(\phi_{2b}-\phi_{2a})f_{11}^b\Big((\phi_{1b}-\phi_{1a})(\chi_{12}+\frac{1}{\phi_{3a}}) - \frac{1}{2v_2}(1-\frac{\phi_{2b}}{\phi_{2a}})\Big), \tag{48}$$

$$f_a = 1, |\mathbf{M}'|_{\chi_{22}} = (\phi_{1b}-\phi_{2b})f_{11}^b\frac{(\phi_{2b}-\phi_{2a})^2}{2v_2\phi_{2b}}. \tag{49}$$

These expressions are first-order infinitesimals. Since $|\mathbf{M}'|_{\chi_{22}}$ is a linear function of $f_a$, it must be a first-order infinitesimal for any $f_a$, so $(\partial\phi_{1a}/\partial\chi_{22})|_{\bar{\phi}_2\to 0}$ must be a second-order infinitesimal for any $f_a$.

##### D3. $(\partial\phi_{1a}/\partial\bar{\phi}_2)|_{\bar{\phi}_2\to 0}$

Let $\bar{\phi}_2 \to 0$ in Eq. (27), we get

$$|\mathbf{M}'|_{\bar{\phi}_2} = (\phi_{1b}-\phi_{1a})f_{11}^{b}\Big((\phi_{1a}-\phi_{1b})\Big(\frac{1}{\phi_{3a}}+\chi_{12}\Big)+\frac{1}{v_2}\Big(1-\frac{\phi_{2b}}{\phi_{2a}}\Big)\Big)\frac{1}{v_2\phi_{2b}}. \tag{50}$$

Substituting the expressions of $\phi_{2b}/\phi_{2a}$ and $\phi_{2a}/\bar{\phi}_2$ into this equation, we get

$$\begin{aligned}\frac{\partial\phi_{1a}}{\partial\bar{\phi}_2}\Big|_{\bar{\phi}_2\to 0} &= \frac{|\mathbf{M}'|_{\bar{\phi}_2}}{|\mathbf{M}|} = \frac{\frac{1}{v_2}\Big(1-\frac{\phi_{2b}}{\phi_{2a}}\Big)-(\phi_{1b}-\phi_{1a})\Big(\frac{1}{\phi_{3a}}+\chi_{12}\Big)}{(\phi_{1b}-\phi_{1a})f_{11}^{a}}\frac{\phi_{2a}}{\bar{\phi}_2}\\ &= \frac{\frac{1}{v_2}(1-e^{-v_2(\chi_{12}(\phi_{1b}-\phi_{1a})-\ln\frac{\phi_{3b}}{\phi_{3a}})})-(\phi_{1b}-\phi_{1a})\Big(\frac{1}{\phi_{3a}}+\chi_{12}\Big)}{(\phi_{1b}-\phi_{1a})f_{11}^{a}(f_a+(1-f_a)e^{-v_2(\chi_{12}(\phi_{1b}-\phi_{1a})-\ln\frac{\phi_{3b}}{\phi_{3a}})})}.\end{aligned} \tag{51}$$

We numerically verify this equation for $v_1 = 100, \chi_{11} = -1.3, f_a = 1$ (Fig. S1).

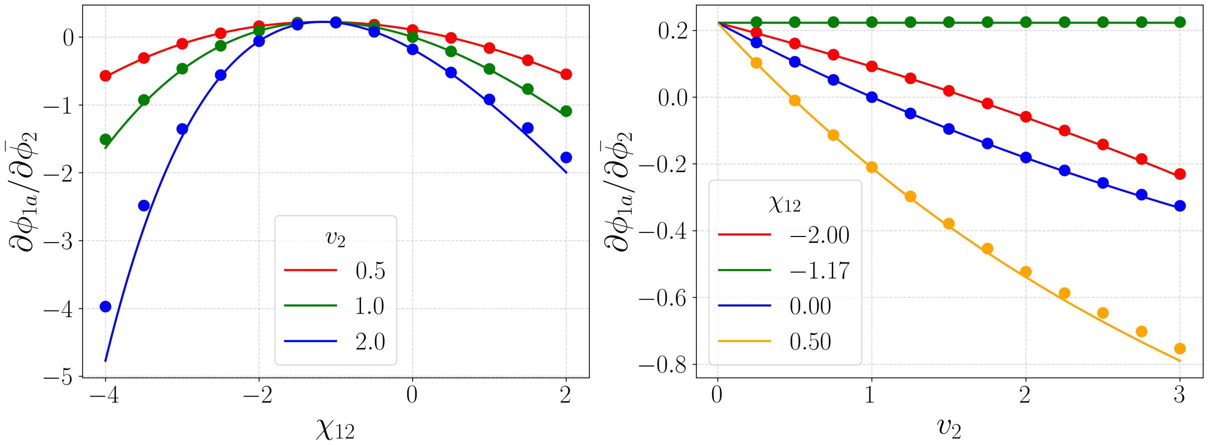


FIG. S1. Comparison of the analytic and numeric values of $(\partial\phi_{1a}/\partial\bar{\phi}_2)|_{\bar{\phi}_2\to 0}$. Analytic values are calculated from Eq. (51) for $v_1 = 100, \chi_{11} = -1.3, f_a = 1$. Numeric values are calculated for $v_1 = 100, \chi_{11} = -1.3$, between $\bar{\phi}_2 = 0$ and $\bar{\phi}_2 = 0.001$.

Here, we consider five special cases.

1) $v_2 \to \infty$. In this case,

$$\frac{\partial\phi_{1a}}{\partial\bar{\phi}_2}\Big|_{\bar{\phi}_2\to 0} = \begin{cases} -\dfrac{\chi_{12}+\frac{1}{\phi_{3a}}}{f_{11}^{a}f_a}, & \chi_{12} > \dfrac{\ln\frac{\phi_{3b}}{\phi_{3a}}}{\phi_{1b}-\phi_{1a}}\\[2ex] -\dfrac{1}{v_2(\phi_{1b}-\phi_{1a})f_{11}^{a}(1-f_a)} \to 0, & \chi_{12} < \dfrac{\ln\frac{\phi_{3b}}{\phi_{3a}}}{\phi_{1b}-\phi_{1a}}.\end{cases} \tag{52}$$

The function exhibits a step change at $\chi_{12}^{c} \equiv \ln(\phi_{3b}/\phi_{3a})/(\phi_{1b}-\phi_{1a})$.

2) $v_2 = 1$. In this case, $\chi_{12} = 0$ is a zero of Eq. (51). Another zero satisfies the following condition:

$$\frac{\phi_{3b}}{\phi_{3a}} = \frac{\chi_{12}(\phi_{1b}-\phi_{1a})}{1-e^{-\chi_{12}(\phi_{1b}-\phi_{1a})}}. \tag{53}$$

3) $v_2 \to 0$. In this case, we consider the zero-level curve of $(\partial\phi_{1a}/\partial\bar{\phi}_2)|_{\bar{\phi}_2\to 0}$. Let $t = v_2\chi_{12}(\phi_{1b}-\phi_{1a}), b = \phi_{3b}/\phi_{3a}$, the zero-level curve is

$$1 - b^{v_2}e^{-t} = v_2(1-b)+t. \tag{54}$$

Expanding the exponential term to the second order, we get

$$1 - b^{v_2}\Big(1 - t + \frac{1}{2}t^2\Big) \approx v_2(1-b) + t. \tag{55}$$

We solve this equation to get

$$t \approx 1 - b^{-v_2} \pm \sqrt{b^{-2v_2} - 1 - 2v_2(1-b)b^{-v_2}}. \tag{56}$$

Performing the following expansions

$$b^{-v_2} \approx 1 - v_2 \ln b, b^{-2v_2} \approx 1 - 2v_2 \ln b, \tag{57}$$

we get

$$t \approx v_2 \ln b \pm \sqrt{-2v_2 \ln b - 2v_2(1-b)}. \tag{58}$$

Dividing both sides by $v_2$, we get

$$\Big(\frac{t}{v_2} - \ln b\Big)^2 \approx \frac{2}{v_2}(b - 1 - \ln b). \tag{59}$$

Plugging in the expressions of $t$ and $b$, we obtain

$$\frac{1}{v_2} \approx \frac{\Big(\chi_{12}(\phi_{1b} - \phi_{1a}) - \ln\frac{\phi_{3b}}{\phi_{3a}}\Big)^2}{2\Big(\frac{\phi_{3b}}{\phi_{3a}} - 1 - \ln\frac{\phi_{3b}}{\phi_{3a}}\Big)} = \frac{(\phi_{1b} - \phi_{1a})^2}{2\Big(\frac{\phi_{3b}}{\phi_{3a}} - 1 - \ln\frac{\phi_{3b}}{\phi_{3a}}\Big)}(\chi_{12} - \chi_{12}^c)^2. \tag{60}$$

We highlight that the above equation shows that the zero-level curve of Eq. (51) has a shape close to a parabola. This approximation conforms better when $v_2$ is small and requires $t = v_2\chi_{12}(\phi_{1b} - \phi_{1a})$ and $v_2 \ln b$ to be small. Apart from letting $v_2 \to 0$, if we let $\chi_{11} \to \chi_{11}^c$, then $\phi_{1b} - \phi_{1a}$ and $\ln b$ will both tend to zero, so we can still perform these expansions.

4) $\chi_{12} = 0$. Since the expression is independent of $\chi_{22}$, the conclusion is consistent with the discussion in section C3, i.e., when $v_2 > 1$, $(\partial\phi_{1a}/\partial\bar{\phi}_2)|_{\bar{\phi}_2\to 0} > 0$; when $v_2 = 1$, $(\partial\phi_{1a}/\partial\bar{\phi}_2)|_{\bar{\phi}_2\to 0} = 0$; when $v_2 < 1$, $(\partial\phi_{1a}/\partial\bar{\phi}_2)|_{\bar{\phi}_2\to 0} < 0$.

5) $\chi_{12} = \chi_{12}^c$. At this point,

$$\frac{\partial\phi_{1a}}{\partial\bar{\phi}_2}\bigg|_{\bar{\phi}_2\to 0} = -\frac{\frac{1}{\phi_{3a}} + \chi_{12}^c}{f_{11}^a}, \tag{61}$$

which is a positive value independent of $v_2$.

#### E. PROPERTIES NEAR THE CRITICAL POINT

##### E1. Landau-Ginzburg theory for binary systems

The Flory-Huggins free energy of a binary system is

$$f(\phi_1) = \frac{1}{v_1}\phi_1 \ln\phi_1 + (1-\phi_1)\ln(1-\phi_1) + \frac{1}{2}\chi_{11}\phi_1^2. \tag{62}$$

At the critical point,

$$\chi_{11}^c = -\frac{(\sqrt{v_1}+1)^2}{v_1}, \quad \phi_1^c = \frac{1}{\sqrt{v_1}+1}. \tag{63}$$

The free energy expanded near the critical point is ($\Delta\phi_1 \equiv \phi_1 - \phi_1^c$, $\Delta\chi_{11} \equiv \chi_{11}^c - \chi_{11}$)

$$f = f^c + \mu_1^c\Delta\phi_1 - \frac{1}{2}(\phi_1^c)^2\Delta\chi_{11} - \phi_1^c\Delta\phi_1\Delta\chi_{11} - \frac{1}{2}(\Delta\phi_1)^2\Delta\chi_{11} + \frac{(\sqrt{v_1}+1)^4}{12\sqrt{v_1^3}}(\Delta\phi_1)^4 + \text{higher order terms}. \tag{64}$$

The first four terms are linear in $\Delta\phi_1$ and do not substantially affect phase separation, so we ignore them:

$$f' = -\frac{1}{2}\Delta\chi_{11}(\Delta\phi_1)^2 + \frac{(\sqrt{v_1}+1)^4}{12\sqrt{v_1^3}}(\Delta\phi_1)^4 + \text{higher order terms.} \tag{65}$$

This is the Landau-Ginzburg expansion of the free energy. It is an even function, and its two minima correspond to the two phases:

$$\varphi \equiv -\Delta\phi_{1a} = \Delta\phi_{1b} = \sqrt{\frac{3\sqrt{v_1^3}}{(\sqrt{v_1}+1)^4}\Delta\chi_{11}} \propto (\Delta\chi_{11})^{\frac{1}{2}}. \tag{66}$$

Here we want to emphasize the condition when this approximation holds, which is $\varphi \ll \phi_1^c$. This condition is satisfied when $\Delta\chi_{11} \ll 1/\sqrt{v_1}$:

$$\sqrt{\frac{3\sqrt{v_1^3}}{(\sqrt{v_1}+1)^4}\Delta\chi_{11}} \ll \frac{1}{\sqrt{v_1}+1} \Rightarrow \Delta\chi_{11} \ll \frac{(\sqrt{v_1}+1)^2}{\sqrt{v_1^3}} \approx \frac{1}{\sqrt{v_1}}. \tag{67}$$

Here, we use the condition $v_1 \gg 1$ in the last approximation of the above equation.

##### E2. Scaling near critical point and critical hypersensitivity

We first look at $\chi_{12}^c$:

$$\chi_{12}^c = \frac{\ln\left(\frac{\phi_{3b}}{\phi_{3a}}\right)}{\phi_{1b}-\phi_{1a}} = \frac{\ln\left(\frac{1-\phi_1^c-\varphi}{1-\phi_1^c+\varphi}\right)}{2\varphi} = -\frac{1}{1-\phi_1^c} - \frac{\varphi^2}{3(1-\phi_1^c)^3} + o(\varphi^3). \tag{68}$$

Define the critical value of $\chi_{12}^c$ as follows:

$$\chi_{12}^{cc} = \lim_{\chi_{11}\to\chi_{11}^c} \chi_{12}^c = -\frac{1}{1-\phi_1^c} = -1 - \frac{1}{\sqrt{v_1}}. \tag{69}$$

We have

$$\chi_{12}^c - \chi_{12}^{cc} \approx -\frac{\varphi^2}{3(1-\phi_1^c)^3}. \tag{70}$$

Next, we perform Taylor expansion to the terms in Eq. (51) and introduce $\Delta\chi_{12} \equiv \chi_{12} - \chi_{12}^c$:

$$\chi_{12}(\phi_{1b}-\phi_{1a}) - \ln\left(\frac{\phi_{3b}}{\phi_{3a}}\right) = 2\varphi(\chi_{12}-\chi_{12}^c) = 2\varphi\Delta\chi_{12}, \tag{71}$$

$$\frac{1}{\phi_{3a}} + \chi_{12} = \Delta\chi_{12} + \frac{1}{1-\phi_1^c+\varphi} + \chi_{12}^c = \Delta\chi_{12} - \frac{\varphi}{(1-\phi_1^c)^2} + o(\varphi), \tag{72}$$

$$\begin{aligned}
f_{11}^a &= \frac{1}{v_1\phi_{1a}} + \frac{1}{\phi_{3a}} + \chi_{11} = \frac{1}{v_1(\phi_1^c-\varphi)} + \frac{1}{1-\phi_1^c+\varphi} + \chi_{11} \\
&= \frac{1}{v_1\phi_1^c}\left(1 + \frac{\varphi}{\phi_1^c} + \frac{\varphi^2}{(\phi_1^c)^2}\right) + \frac{1}{1-\phi_1^c}\left(1 - \frac{\varphi}{1-\phi_1^c} + \frac{\varphi^2}{(1-\phi_1^c)^2}\right) + \chi_{11} + o(\varphi^2) \\
&= (\chi_{11}-\chi_{11}^c) + \left(\frac{1}{v_1(\phi_1^c)^3} + \frac{1}{(1-\phi_1^c)^3}\right)\varphi^2 + o(\varphi^2) \\
&= -\frac{(\sqrt{v_1}+1)^4}{3\sqrt{v_1^3}}\varphi^2 + \frac{(\sqrt{v_1}+1)^4}{\sqrt{v_1^3}}\varphi^2 + o(\varphi^2) = \frac{2(\sqrt{v_1}+1)^4}{3\sqrt{v_1^3}}\varphi^2 + o(\varphi^2).
\end{aligned} \tag{73}$$

Substituting these expansions into Eq. (51), we obtain

$$\frac{\partial \phi_{1a}}{\partial \bar{\phi}_2}\bigg|_{\bar{\phi}_2\to 0} = \frac{\frac{1}{v_2}(1-e^{-2v_2\Delta\chi_{12}\varphi+o(\varphi^2)}) - 2\Delta\chi_{12}\varphi + \frac{2}{(1-\phi_1^c)^2}\varphi^2 + o(\varphi^2)}{2\varphi\Big(\frac{2(\sqrt{v_1}+1)^4}{3\sqrt{v_1^3}}\varphi^2 + o(\varphi^2)\Big)(f_a + (1-f_a)e^{-2v_2\Delta\chi_{12}\varphi+o(\varphi^2)})}. \tag{74}$$

According to the limit of the exponent, we discuss two cases.

1) $v_2\varphi \ll 1$, which means the absolute value of the exponent tends to 0. This is true in most cases as long as $v_2$ is not very large, because $\varphi$ is very small near the critical point. In this case, we expand the exponential function to get

$$\frac{\partial \phi_{1a}}{\partial \bar{\phi}_2}\bigg|_{\bar{\phi}_2\to 0} \approx \Big(\frac{1}{(1-\phi_1^c)^2} - v_2\Delta\chi_{12}^2\Big)\frac{3\sqrt{v_1^3}}{2(\sqrt{v_1}+1)^4}\varphi^{-1} \propto \varphi^{-1}. \tag{75}$$

Therefore, as the system approaches the critical point, $(\partial\phi_{1a}/\partial\bar{\phi}_2)|_{\bar{\phi}_2\to 0}$ tends to infinity, which means that the system exhibits critical hypersensitivity when a macromolecule is added.

From this equation, we get the following zero-level curve:

$$v_2\Delta\chi_{12}^2 \approx \frac{1}{(1-\phi_1^c)^2}. \tag{76}$$

This expression can also be obtained by taking the limit of Eq. (60):

$$\begin{aligned} v_2(\chi_{12}-\chi_{12}^c)^2 &\approx \frac{2(\frac{\phi_{3b}}{\phi_{3a}} - 1 - \ln\frac{\phi_{3b}}{\phi_{3a}})}{(\phi_{1b}-\phi_{1a})^2} \approx \frac{2}{(2\varphi)^2}\Big(\frac{1-\phi_1^c-\varphi}{1-\phi_1^c+\varphi} - 1 - \ln\frac{1-\phi_1^c-\varphi}{1-\phi_1^c+\varphi}\Big) \\ &\approx \frac{1}{2\varphi^2}\Big(1 - \frac{2\varphi}{1-\phi_1^c} + \frac{2\varphi^2}{(1-\phi_1^c)^2} - 1 + \frac{2\varphi}{1-\phi_1^c}\Big) = \frac{1}{(1-\phi_1^c)^2} \end{aligned} \tag{77}$$

2) $v_2\varphi \gg 1$, which means the absolute value of the exponent tends to infinity, corresponding to very large $v_2$. In this case, the sign of the exponent depends on $\chi_{12}$. When $\chi_{12} < \chi_{12}^c$, the exponential function tends to infinity. Ignoring other terms that are not exponential, we obtain

$$\frac{\partial \phi_{1a}}{\partial \bar{\phi}_2}\bigg|_{\bar{\phi}_2\to 0} \approx -\frac{3\sqrt{v_1^3}}{4(\sqrt{v_1}+1)^4}\frac{1}{(1-f_a)v_2\varphi^3} \propto \frac{\varphi^{-3}}{v_2}. \tag{78}$$

When $\chi_{12} > \chi_{12}^c$, the exponential function tends to 0 and can be directly ignored, yielding

$$\frac{\partial \phi_{1a}}{\partial \bar{\phi}_2}\bigg|_{\bar{\phi}_2\to 0} \approx -\frac{3\sqrt{v_1^3}}{2(\sqrt{v_1}+1)^4}\frac{\Delta\chi_{12}}{f_a\varphi^2} \propto \varphi^{-2}. \tag{79}$$

##### E3. Conclusions of Landau-Ginzburg theory for $\partial\phi_{1b}/\partial\bar{\phi}_2$

Similar to Eq. (27), but this time we use $\vec{\beta}_{\bar{\phi}_2}$ to replace the second column in matrix $\mathbf{M}$ to get $\mathbf{M}'$, from which we get

$$\frac{\partial \phi_{1b}}{\partial \bar{\phi}_2}\bigg|_{\bar{\phi}_2\to 0} = \frac{\frac{1}{v_2}(e^{v_2(\chi_{12}(\phi_{1b}-\phi_{1a}) - \ln\frac{\phi_{3b}}{\phi_{3a}})} - 1) - (\phi_{1b}-\phi_{1a})\Big(\frac{1}{\phi_{3b}} + \chi_{12}\Big)}{(\phi_{1b}-\phi_{1a})f_{11}^b(f_a e^{v_2(\chi_{12}(\phi_{1b}-\phi_{1a}) - \ln\frac{\phi_{3b}}{\phi_{3a}})} + 1 - f_a)}. \tag{80}$$

Similarly, we find the expression of $(\partial\phi_{1b}/\partial\bar{\phi}_2)|_{\bar{\phi}_2\to 0}$ near the critical point as

$$\frac{\partial \phi_{1b}}{\partial \bar{\phi}_2}\bigg|_{\bar{\phi}_2\to 0} = \frac{\frac{1}{v_2}(e^{2v_2\Delta\chi_{12}\varphi+o(\varphi^2)} - 1) - 2\Delta\chi_{12}\varphi - \frac{2}{(1-\phi_1^c)^2}\varphi^2 + o(\varphi^2)}{2\varphi\Big(\frac{2(\sqrt{v_1}+1)^4}{3\sqrt{v_1^3}}\varphi^2 + o(\varphi^2)\Big)(f_a e^{2v_2\Delta\chi_{12}\varphi+o(\varphi^2)} + 1 - f_a)}. \tag{81}$$

Based on the limit of the exponential function, we discuss two cases.

1) $v_2\varphi \ll 1$, which means the absolute value of the exponent tends to 0.

$$\frac{\partial \phi_{1b}}{\partial \bar{\phi}_2}\bigg|_{\bar{\phi}_2\to 0} \approx \Big(v_2\Delta\chi_{12}^2 - \frac{1}{(1-\phi_1^c)^2}\Big)\frac{3\sqrt{v_1^3}}{2(\sqrt{v_1}+1)^4}\varphi^{-1} \propto \varphi^{-1}, \tag{82}$$

which is exactly the opposite of $(\partial\phi_{1a}/\partial\bar{\phi}_2)|_{\bar{\phi}_2\to 0}$. The limits of their zero-level curves are the same, both being $v_2\Delta\chi_{12}^2 = 1/(1-\phi_1^c)^2$.

2) $v_2\varphi \gg 1$. When $\chi_{12} < \chi_{12}^c$, the exponential function tends to 0.

$$\frac{\partial \phi_{1b}}{\partial \bar{\phi}_2}\bigg|_{\bar{\phi}_2\to 0} \approx -\frac{3\sqrt{v_1^3}}{2(\sqrt{v_1}+1)^4}\frac{\Delta\chi_{12}}{(1-f_a)\varphi^2} \propto \varphi^{-2}. \tag{83}$$

When $\chi_{12} > \chi_{12}^c$, the exponential function tends to infinity.

$$\frac{\partial \phi_{1b}}{\partial \bar{\phi}_2}\bigg|_{\bar{\phi}_2\to 0} \approx \frac{3\sqrt{v_1^3}}{4(\sqrt{v_1}+1)^4}\frac{1}{f_a v_2\varphi^3} \propto \frac{\varphi^{-3}}{v_2}. \tag{84}$$

#### F. BIOLOGICALLY RELEVANT SCENARIO: LARGE $v_1$

When $v_1$ is large, as $\Delta\chi_{11}$ gradually increases, the system can be divided into three scenarios. Fig. S2 shows the free energy as a function of $\phi_1$ and its tangent line in these three scenarios. The first scenario corresponds to the two phases having similar volume fractions, which can be treated by the Landau-Ginzburg expansion discussed in section E; the second scenario corresponds to the dilute phase volume fraction approaching 0, while the dense phase volume fraction is still small; the third scenario corresponds to the dilute phase volume fraction approaching 0, while the dense phase volume fraction approaches 1. We have discussed the first scenario, and here we will focus on the second scenario, which is more biologically relevant.

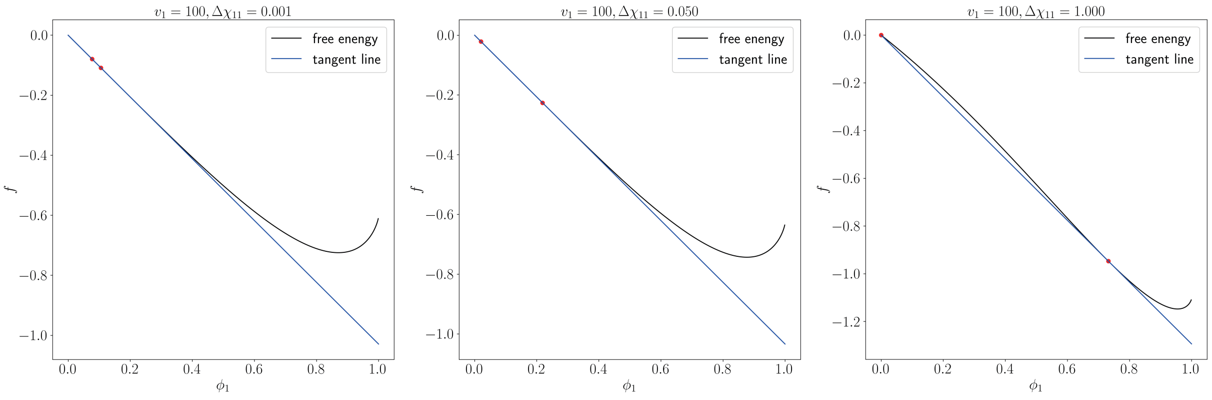


FIG. S2. Three scenarios under different $\Delta\chi_{11}$ when $v_1$ is large

When $v_1$ is large, $\phi_1^c$ is small. The biologically relevant scenario discussed here corresponds to $\phi_{1a}$ being several orders of magnitude smaller than $\phi_1^c$, while $\phi_{1b}$ is greater than $\phi_1^c$ but still much smaller than one. In other words, $0 < \phi_{1a} \ll \phi_1^c < \phi_{1b} \ll 1$. As we show in the following, we require $v_1$ to be large enough for the condition $0 < \phi_{1a} \ll \phi_1^c < \phi_{1b} \ll 1$ to hold.

When $0 < \phi_{1a} \ll \phi_1^c < \phi_{1b} \ll 1$, we can ignore all terms containing $\phi_{1a}$ in Eq. (36) except for $\ln\phi_{1a}$. We get

$$\Big(\frac{1}{v_1} - 1\Big)\phi_{1b} - \ln(1-\phi_{1b}) + \frac{1}{2}\chi_{11}\phi_{1b}^2 = 0 \tag{85}$$

and

$$\frac{1}{v_1}\ln\phi_{1a} = \frac{1}{v_1}\ln\phi_{1b} - \ln(1-\phi_{1b}) + \chi_{11}\phi_{1b}. \tag{86}$$

When $v_1$ is large, $\chi_{11}^c \to -1$, so we define $\Delta\chi_{11} \equiv -\chi_{11} - 1 > 0$.First, we expand Eq. (85) to the third order of $\phi_{1b}$:

$$\Big(\frac{1}{v_1} - 1\Big)\phi_{1b} + \phi_{1b} + \frac{1}{2}\phi_{1b}^2 + \frac{1}{3}\phi_{1b}^3 + \frac{1}{2}\chi_{11}\phi_{1b}^2 = 0. \tag{87}$$

Since $\phi_{1b} \neq 0$,

$$\phi_{1b}^2 - \frac{3}{2}\Delta\chi_{11}\phi_{1b} + \frac{3}{v_1} = 0. \tag{88}$$

Solving this equation by quadratic formula and expanding to the first order of $1/v_1$, we get

$$\phi_{1b} = \frac{3}{4}\Delta\chi_{11}\Big(1 + \sqrt{1 - \frac{16}{3v_1\Delta\chi_{11}^2}}\Big) \approx \frac{3}{2}\Delta\chi_{11}\Big(1 - \frac{4}{3v_1\Delta\chi_{11}^2}\Big). \tag{89}$$

Plugging into Eq. (86), we obtain

$$\begin{aligned}
\phi_{1a} &= \phi_{1b}\exp(v_1(\chi_{11}\phi_{1b} - \ln(1-\phi_{1b}))) \\
&= \phi_{1b}\exp\Big(v_1\Big(\chi_{11}\phi_{1b} - \Big(\frac{1}{v_1} - 1\Big)\phi_{1b} - \frac{1}{2}\chi_{11}\phi_{1b}^2\Big)\Big) \\
&\approx \phi_{1b}\exp\Big(\frac{3}{2}v_1\Delta\chi_{11}\Big(1 - \frac{4}{3v_1\Delta\chi_{11}^2}\Big)\Big(-\Delta\chi_{11} - \frac{1}{v_1} - \frac{1}{2}(-\Delta\chi_{11} - 1)\Big(\frac{3}{2}\Delta\chi_{11}\Big(1 - \frac{4}{3v_1\Delta\chi_{11}^2}\Big)\Big)\Big)\Big) \\
&\approx \phi_{1b}\exp\Big(\frac{3}{2}\Big(1 - \frac{4}{3v_1\Delta\chi_{11}^2}\Big)\Big(-v_1\Delta\chi_{11}^2 + \frac{3}{4}v_1\Delta\chi_{11}^2\Big(1 - \frac{4}{3v_1\Delta\chi_{11}^2}\Big)\Big)\Big) \\
&= \phi_{1b}\exp\Big(-\frac{3}{2}\Big(v_1\Delta\chi_{11}^2 - \frac{4}{3}\Big)\Big(\frac{1}{4} + \frac{1}{v_1\Delta\chi_{11}^2}\Big)\Big) \\
&\approx \phi_{1b}\exp\Big(-\frac{3}{8}v_1\Delta\chi_{11}^2 - 1\Big).
\end{aligned} \tag{90}$$

In deriving the above equation, we replace $\ln(1-\phi_{1b})$ by $(1/v_1 - 1)\phi_{1b} + \chi_{11}\phi_{1b}^2/2$ using Eq. (85) and use the two approximations $v_1\Delta\chi_{11}^2 \gg 1$ and $\Delta\chi_{11} \ll 1$. We note that given the condition $0 < \phi_{1a} \ll \phi_1^c < \phi_{1b} \ll 1$, these two approximations must hold. This is because $\phi_{1a}$ needs to be much smaller than $\phi_{1b}$, so the absolute value of the exponent in the last line of Eq. (90) must be much greater than 1, which means that

$$v_1\Delta\chi_{11}^2 \gg 1 \Rightarrow \Delta\chi_{11} \gg \frac{1}{\sqrt{v_1}}. \tag{91}$$

Meanwhile, $\phi_{1b} \ll 1$ requires that $\Delta\chi_{11} \ll 1$. Therefore, to satisfy $0 < \phi_{1a} \ll \phi_1^c < \phi_{1b} \ll 1$, we need to have

$$\frac{1}{\sqrt{v_1}} \ll \Delta\chi_{11} \ll 1. \tag{92}$$

Clearly, for the above condition on $\Delta\chi_{11}$ to hold, we must have $v_1 \gg 1$. To summarize, when $\Delta\chi_{11} \ll 1/\sqrt{v_1}$, i.e. the system is very close to the critical point, we can use Landau-Ginzburg expansion; when $1/\sqrt{v_1} \ll \Delta\chi_{11} \ll 1$, i.e. the system is moderately close to the critical point, we can use the approximations here.

Now that we have $v_1\Delta\chi_{11}^2 \gg 1$, we can further approximate as follows:

$$\phi_{1b} \approx \frac{3}{2}\Delta\chi_{11}, \phi_{1a} \approx \frac{3}{2e}\Delta\chi_{11}\exp\Big(-\frac{3}{8}v_1\Delta\chi_{11}^2\Big), \tag{93}$$

$$\begin{aligned}
\ln\frac{\phi_{2b}}{\phi_{2a}} &= -v_2\Big(\chi_{12}(\phi_{1b} - \phi_{1a}) - \ln\frac{1-\phi_{1b}}{1-\phi_{1a}}\Big) \\
&\approx -v_2(\chi_{12}\phi_{1b} - \ln(1-\phi_{1b})) \\
&\approx -v_2\Big(\chi_{12}\phi_{1b} + \phi_{1b} + \frac{1}{2}\phi_{1b}^2\Big) \\
&\approx -v_2\Big(\frac{3}{2}\Delta\chi_{11}\Big)\Big(\Delta\chi_{12} + \frac{1}{2}\Big(\frac{3}{2}\Delta\chi_{11}\Big)\Big) \\
&= -\frac{3}{2}v_2\Delta\chi_{11}\Big(\Delta\chi_{12} + \frac{3}{4}\Delta\chi_{11}\Big) \\
&\approx -\frac{3}{2}v_2\Delta\chi_{11}\Delta\chi_{12},
\end{aligned} \tag{94}$$

where $\phi_{2b}/\phi_{2a}$ is the partition coefficient of the macromolecule between two phases. We numerically test the approximation of $\ln(\phi_{2b}/\phi_{2a})$ in Fig. S3.

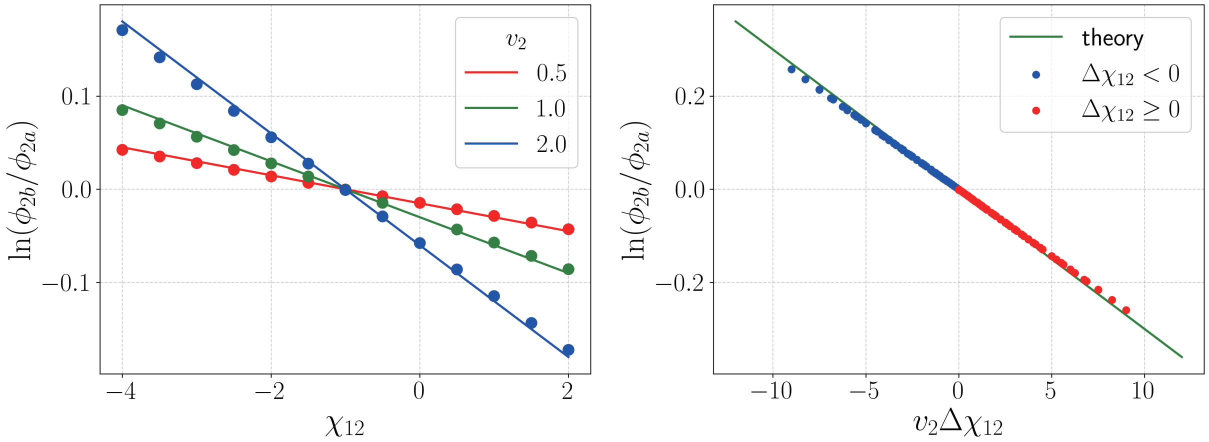


FIG. S3. The partition coefficient of the macromolecule between the two phases. The data points are from numerical calculations, and the solid lines are theoretical predictions [the last line of Eq. (94)]. In this figure, $v_1 = 10^5$, $\chi_{11} = -1.02$, $f_a = 0.9$, and the data points are calculated between $\bar{\phi}_2 = 0$ and $\bar{\phi}_2 = 10^{-5}$.

We first discuss $(\partial\phi_{1a}/\partial\bar{\phi}_2)|_{\bar{\phi}_2\to0}$. When $v_1$ is large, $\chi_{12}^{cc} \to -1$, so we define $\Delta\chi_{12} \equiv \chi_{12} + 1$. We substitute the approximations of $\phi_{1a}$ and $\phi_{1b}$ in Eq. (93) into Eq. (51) to get

$$\begin{aligned}
\left.\frac{\partial\phi_{1a}}{\partial\bar{\phi}_2}\right|_{\bar{\phi}_2\to0} &= \frac{\frac{1}{v_2}\left(1-e^{-(v_2(\chi_{12}(\phi_{1b}-\phi_{1a})-\ln\frac{\phi_{3b}}{\phi_{3a}}))}\right)-(\phi_{1b}-\phi_{1a})\left(\frac{1}{\phi_{3a}}+\chi_{12}\right)}{(\phi_{1b}-\phi_{1a})f_{11}^{a}\left(f_a+(1-f_a)e^{-v_2(\chi_{12}(\phi_{1b}-\phi_{1a})-\ln\frac{\phi_{3b}}{\phi_{3a}})}\right)} \\
&\approx \frac{\frac{1}{v_2}\left(1-e^{-v_2(\Delta\chi_{12}\phi_{1b}+\frac{1}{2}\phi_{1b}^2)}\right)-\phi_{1b}\Delta\chi_{12}}{f_a+(1-f_a)e^{-v_2\Delta\chi_{12}\phi_{1b}}}\frac{v_1\phi_{1a}}{\phi_{1b}} \\
&\approx \frac{1}{2}(1-v_2\Delta\chi_{12}^2)\phi_{1b}^2\frac{v_1\phi_{1a}}{\phi_{1b}} \\
&\approx \frac{9}{8e}(1-v_2\Delta\chi_{12}^2)v_1\Delta\chi_{11}^2e^{-\frac{3}{8}v_1\Delta\chi_{11}^2}.
\end{aligned} \tag{95}$$

Here, the second step of approximation requires $v_2$ not to be too large. The zero-level curve is still $v_2\Delta\chi_{12}^2 = 1$ in this scenario, which is consistent with Eq. (76) when $\phi_1^c \to 0$.

Then, we discuss $(\partial\phi_{1b}/\partial\bar{\phi}_2)|_{\bar{\phi}_2\to0}$:

$$\begin{aligned}
\left.\frac{\partial\phi_{1b}}{\partial\bar{\phi}_2}\right|_{\bar{\phi}_2\to0} &= \frac{\frac{1}{v_2}\left(e^{v_2(\chi_{12}(\phi_{1b}-\phi_{1a})-\ln\frac{\phi_{3b}}{\phi_{3a}})}-1\right)-(\phi_{1b}-\phi_{1a})\left(\frac{1}{\phi_{3b}}+\chi_{12}\right)}{(\phi_{1b}-\phi_{1a})f_{11}^{b}\left(f_ae^{v_2(\chi_{12}(\phi_{1b}-\phi_{1a})-\ln\frac{\phi_{3b}}{\phi_{3a}})}+1-f_a\right)} \\
&\approx \frac{\frac{1}{v_2}\left(e^{v_2(\Delta\chi_{12}\phi_{1b}+\frac{1}{2}\phi_{1b}^2)}-1\right)-\phi_{1b}(\Delta\chi_{12}+\phi_{1b})}{\frac{1}{3}\phi_{1b}^2\left(f_ae^{v_2\Delta\chi_{12}\phi_{1b}}+1-f_a\right)} \\
&\approx \frac{\frac{1}{2}\phi_{1b}^2(v_2\Delta\chi_{12}^2-1)}{\frac{1}{3}\phi_{1b}^2} \approx \frac{3}{2}(v_2\Delta\chi_{12}^2-1).
\end{aligned} \tag{96}$$

The result is independent of $\Delta\chi_{11}$. The zero-level curve is still $v_2\Delta\chi_{12}^2 = 1$.

#### G. EFFECT OF MACROMOLECULE ON THE VOLUME RATIO: $(\partial f_b/\partial\bar{\phi}_2)|_{\bar{\phi}_2\to 0}$

From conservation of mass, we have

$$(1-f_b)\phi_{1a}+f_b\phi_{1b}=\bar{\phi}_1. \tag{97}$$

Differentiating both sides of the equation with respect to $\bar{\phi}_2$ yields

$$\frac{\partial f_b}{\partial\bar{\phi}_2}=-\frac{(1-f_b)\frac{\partial\phi_{1a}}{\partial\bar{\phi}_2}+f_b\frac{\partial\phi_{1b}}{\partial\bar{\phi}_2}}{\phi_{1b}-\phi_{1a}}. \tag{98}$$

Substituting Eq. (51) and Eq. (80), we get

$$\begin{aligned}\left.\frac{\partial f_b}{\partial\bar{\phi}_2}\right|_{\bar{\phi}_2\to 0}=&-(1-f_b)\frac{\frac{1}{v_2}(1-e^{-(v_2(\chi_{12}(\phi_{1b}-\phi_{1a})-\ln\frac{\phi_{3b}}{\phi_{3a}}))})-(\phi_{1b}-\phi_{1a})\Big(\frac{1}{\phi_{3a}}+\chi_{12}\Big)}{(\phi_{1b}-\phi_{1a})^2f_{11}^a(f_a+(1-f_a)e^{-v_2(\chi_{12}(\phi_{1b}-\phi_{1a})-\ln\frac{\phi_{3b}}{\phi_{3a}})})}\\&-f_b\frac{\frac{1}{v_2}(e^{v_2(\chi_{12}(\phi_{1b}-\phi_{1a})-\ln\frac{\phi_{3b}}{\phi_{3a}})}-1)-(\phi_{1b}-\phi_{1a})\Big(\frac{1}{\phi_{3b}}+\chi_{12}\Big)}{(\phi_{1b}-\phi_{1a})^2f_{11}^b(f_ae^{v_2(\chi_{12}(\phi_{1b}-\phi_{1a})-\ln\frac{\phi_{3b}}{\phi_{3a}})}+1-f_a)}.\end{aligned} \tag{99}$$

This expression contains $f_b$. For different scenarios discussed in previous sections, $(\partial f_b/\partial\bar{\phi}_2)|_{\bar{\phi}_2\to 0}$ can be calculated using Eq. (98). For example, for the biologically relevant scenario discussed in section F ($1/\sqrt{v_1}\ll\Delta\chi_{11}\ll 1$), $(\partial\phi_{1a}/\partial\bar{\phi}_2)\ll(\partial\phi_{1b}/\partial\bar{\phi}_2)$, so we have

$$\frac{\partial f_b}{\partial\bar{\phi}_2}\approx-\frac{f_b}{\phi_{1b}}\frac{\partial\phi_{1b}}{\partial\bar{\phi}_2}\approx-\frac{3f_b}{2\phi_{1b}}(v_2\Delta\chi_{12}^2-1). \tag{100}$$

#### H. EFFECT OF MACROMOLECULE ON MULTI-COMPONENT TWO-PHASE SYSTEM

We generalize our theory to a solution of $n$ protein solutes (labeled from 1 to $n$) and a macromolecule cosolute (labeled $m\equiv n+1$) in solvent $s$. The Flory-Huggins free energy is

$$f=\sum_{i=1}^{n+1}\frac{1}{v_i}\phi_i\ln\phi_i+(1-\sum_{i=1}^{n+1}\phi_i)\ln(1-\sum_{i=1}^{n+1}\phi_i)+\frac{1}{2}\sum_{i=1}^{n+1}\sum_{j=1}^{n+1}\chi_{ij}\phi_i\phi_j. \tag{101}$$

We consider the case when the system separates into two phases, a and b. At equilibrium,

$$\begin{cases}\mu_{ia}-\mu_{ib} &=0,\quad i=1\sim n+1 & \text{(102a)}\\ \Pi_a-\Pi_b &=0 & \text{(102b)}\\ f_a\phi_{ia}+(1-f_a)\phi_{ib}-\bar{\phi}_i &=0,\quad i=1\sim n+1, & \text{(102c)}\end{cases}$$

where

$$\mu_i=\frac{\partial f}{\partial\phi_i},\quad\frac{\partial\mu_i}{\partial\phi_j}\equiv f_{ij}, \tag{103}$$

$$\Pi=-f+\sum_{i=1}^{n+1}\phi_i\mu_i\Rightarrow\frac{\partial\Pi}{\partial\phi_j}=\sum_{i=1}^{n+1}\phi_if_{ij}. \tag{104}$$

Similar to Eq. (8), we consider the linear responses against the addition of macromolecule:

$$\begin{pmatrix} f^a_{11} & ... & f^a_{1(n+1)} & -f^b_{11} & ... & -f^b_{1(n+1)} & 0 \\ ... & ... & ... & ... & ... & ... & ... \\ f^a_{1(n+1)} & ... & f^a_{(n+1)(n+1)} & -f^b_{1(n+1)} & ... & -f^b_{(n+1)(n+1)} & 0 \\ \sum_{i=1}^{n+1}\phi_{ia}f^a_{1i} & ... & \sum_{i=1}^{n+1}\phi_{ia}f^a_{(n+1)i} & -\sum_{i=1}^{n+1}\phi_{ib}f^b_{1i} & ... & -\sum_{i=1}^{n+1}\phi_{ib}f^b_{(n+1)i} & 0 \\ f_a & ... & 0 & 1-f_a & ... & 0 & \phi_{1a}-\phi_{1b} \\ ... & ... & ... & ... & ... & ... & ... \\ 0 & ... & f_a & 0 & ... & 1-f_a & \phi_{(n+1)a}-\phi_{(n+1)b} \end{pmatrix} \begin{pmatrix} \delta\phi_{1a} \\ ... \\ \delta\phi_{(n+1)a} \\ \delta\phi_{1b} \\ ... \\ \delta\phi_{(n+1)b} \\ \delta f_a \end{pmatrix} = \begin{pmatrix} 0 \\ ... \\ 0 \\ 0 \\ ... \\ 0 \\ \delta\bar{\phi}_m \end{pmatrix}. \tag{105}$$

We define the Hessian matrix and vector $\phi$ as follows:

$$\mathbf{H} = \{f_{ij}\}_{1\le i,j\le n+1}, \quad \phi = \{\phi_i\}_{1\le i\le n+1}. \tag{106}$$

This allows us to rewrite the equation in block matrices, where $\mathbf{I_m}$ is the identity matrix $(m\times m)$ and $\mathbf{e}_m$ is the unit vector in the m-th dimension $(m\times 1)$:

$$\begin{pmatrix} \mathbf{H}^a & -\mathbf{H}^b & \mathbf{0} \\ \phi_a^{\mathrm{T}}\mathbf{H}^a & -\phi_b^{\mathrm{T}}\mathbf{H}^b & 0 \\ f_a\mathbf{I_m} & (1-f_a)\mathbf{I_m} & \phi_a-\phi_b \end{pmatrix} \begin{pmatrix} \dfrac{\partial\phi_a}{\partial\bar{\phi}_m} \\ \dfrac{\partial\phi_b}{\partial\bar{\phi}_m} \\ \dfrac{\partial f_a}{\partial\bar{\phi}_m} \end{pmatrix} = \begin{pmatrix} \mathbf{0} \\ 0 \\ \mathbf{e}_m \end{pmatrix}. \tag{107}$$

##### H1. The case with a major phase dominating in volume, $f_a = 1$

First, we consider the case: $f_a = 1$. In this case, the system is right at the boundary in the parameter space between the single-phase region and the two-phase region, i.e., at the edge of forming condensates. In this case,

$$\begin{aligned} |\mathbf{M}| &= \begin{vmatrix} \mathbf{H}^a & -\mathbf{H}^b & \mathbf{0} \\ \phi_a^{\mathrm{T}}\mathbf{H}^a & -\phi_b^{\mathrm{T}}\mathbf{H}^b & 0 \\ \mathbf{I_m} & \mathbf{0} & \phi_a-\phi_b \end{vmatrix} = \begin{vmatrix} \mathbf{H}^a & -\mathbf{H}^b & \mathbf{0} \\ (\phi_a^{\mathrm{T}}-\phi_b^{\mathrm{T}})\mathbf{H}^a & \mathbf{0} & 0 \\ \mathbf{I_m} & \mathbf{0} & \phi_a-\phi_b \end{vmatrix} = |\mathbf{H}^b| \begin{vmatrix} (\phi_a^{\mathrm{T}}-\phi_b^{\mathrm{T}})\mathbf{H}^a & 0 \\ \mathbf{I_m} & \phi_a-\phi_b \end{vmatrix} \\ &= |\mathbf{H}^b| \begin{vmatrix} (\phi_a^{\mathrm{T}}-\phi_b^{\mathrm{T}})\mathbf{H}^a & -(\phi_a^{\mathrm{T}}-\phi_b^{\mathrm{T}})\mathbf{H}^a(\phi_a-\phi_b) \\ \mathbf{I_m} & \mathbf{0} \end{vmatrix} = (-1)^n(\phi_a^{\mathrm{T}}-\phi_b^{\mathrm{T}})\mathbf{H}^a(\phi_a-\phi_b)|\mathbf{H}^b|. \end{aligned} \tag{108}$$

We can calculate $|\mathbf{M}'|$ as we do in Eq. (27). We define $\tilde{\mathbf{H}} \equiv \{f_{ij}\}_{1\le i\le n+1, 2\le j\le n+1}$, i.e. remove the first column of $\mathbf{H}$, and $\tilde{\phi} \equiv \{\phi_i\}_{2\le i\le n+1}$. We have

$$\begin{aligned} |\mathbf{M}'| &= \begin{vmatrix} \mathbf{0} & \tilde{\mathbf{H}}^a & -\mathbf{H}^b & \mathbf{0} \\ 0 & \phi_a^{\mathrm{T}}\tilde{\mathbf{H}}^a & -\phi_b^{\mathrm{T}}\mathbf{H}^b & 0 \\ 0 & \mathbf{0} & \mathbf{0} & \phi_{1a}-\phi_{1b} \\ \mathbf{e}_n & \mathbf{I}_n & \mathbf{0} & \tilde{\phi}_a-\tilde{\phi}_b \end{vmatrix} = (-1)^n(\phi_{1a}-\phi_{1b}) \begin{vmatrix} \mathbf{0} & \tilde{\mathbf{H}}^a & -\mathbf{H}^b \\ 0 & \phi_a^{\mathrm{T}}\tilde{\mathbf{H}}^a & -\phi_b^{\mathrm{T}}\mathbf{H}^b \\ \mathbf{e}_n & \mathbf{I}_n & \mathbf{0} \end{vmatrix} \\ &= (-1)^n(\phi_{1a}-\phi_{1b})|\mathbf{H}^b| \begin{vmatrix} 0 & (\phi_a^{\mathrm{T}}-\phi_b^{\mathrm{T}})\tilde{\mathbf{H}}^a \\ \mathbf{e}_n & \mathbf{I}_n \end{vmatrix} = (-1)^{n+1}(\phi_{1a}-\phi_{1b})|\mathbf{H}^b|(\phi_a^{\mathrm{T}}-\phi_b^{\mathrm{T}})\tilde{\mathbf{H}}^a\mathbf{e}_n. \end{aligned} \tag{109}$$

Therefore,

$$\frac{\partial \phi_{1a}}{\partial \bar{\phi}_m} = \frac{|\mathbf{M}'|}{|\mathbf{M}|} = \frac{(\phi_{1b}-\phi_{1a})(\phi_a^{\mathrm{T}}-\phi_b^{\mathrm{T}})\tilde{\mathbf{H}}^a\mathbf{e}_n}{(\phi_a^{\mathrm{T}}-\phi_b^{\mathrm{T}})\mathbf{H}^a(\phi_a-\phi_b)} = \frac{(\phi_{1b}-\phi_{1a})}{(\phi_a^{\mathrm{T}}-\phi_b^{\mathrm{T}})\mathbf{H}^a(\phi_a-\phi_b)}\sum_{i=1}^{n+1}(\phi_{ia}-\phi_{ib})f_{im}^a. \tag{110}$$

Since $\phi_{1b}-\phi_{1a}>0$, and the quadratic form in the denominator is also positive, the sign of $\partial\phi_{1a}/\partial\bar{\phi}_m$ is determined by $\sum_{i=1}^{n+1}(\phi_{ia}-\phi_{ib})f_{im}^a$. For convenience, we denote the sum as $Q$.

We consider the case when $\bar{\phi}_m \to 0$:

$$\begin{aligned} Q \equiv \sum_{i=1}^{n+1}(\phi_{ia}-\phi_{ib})f_{im}^a &= \sum_{i=1}^{n}(\phi_{ia}-\phi_{ib})\Big(\frac{1}{1-\sum_{j=1}^{n+1}\phi_{ja}}+\chi_{im}\Big)+(\phi_{ma}-\phi_{mb})\Big(\frac{1}{v_m\phi_{ma}}+\frac{1}{1-\sum_{j=1}^{n+1}\phi_{ja}}+\chi_{mm}\Big) \\ &\to \sum_{i=1}^{n}(\phi_{ia}-\phi_{ib})\Big(\frac{1}{1-\sum_{j=1}^{n}\phi_{ja}}+\chi_{im}\Big)+\frac{1}{v_m}\Big(1-\frac{\phi_{mb}}{\phi_{ma}}\Big), \end{aligned} \tag{111}$$

where

$$\mu_m^a=\mu_m^b \Rightarrow \ln\Big(\frac{\phi_{mb}}{\phi_{ma}}\Big)=v_m\Big(-\sum_{i=1}^{n}\chi_{im}(\phi_{ib}-\phi_{ia})+\ln\Big(\frac{1-\sum_{i=1}^{n}\phi_{ib}}{1-\sum_{i=1}^{n}\phi_{ia}}\Big)\Big). \tag{112}$$

We assume that the solution is dilute, i.e. $\phi_{ia}\ll 1, \phi_{ib}\ll 1$. Expanding Q to the second order, we get

$$\ln\Big(\frac{\phi_{mb}}{\phi_{ma}}\Big)\approx -v_m\Big(\sum_{i=1}^{n}(\chi_{im}+1)(\phi_{ib}-\phi_{ia})+\frac{1}{2}(\sum_{i=1}^{n}\phi_{ib})^2-\frac{1}{2}(\sum_{i=1}^{n}\phi_{ia})^2\Big), \tag{113}$$

$$\begin{aligned} Q \approx& \sum_{i=1}^{n}(\phi_{ia}-\phi_{ib})(1+\sum_{j=1}^{n}\phi_{ja}+\chi_{im})+\frac{1}{v_m}\Big(1-\exp(-v_m[\sum_{i=1}^{n}(\chi_{im}+1)(\phi_{ib}-\phi_{ia})+\frac{1}{2}(\sum_{i=1}^{n}\phi_{ib})^2-\frac{1}{2}(\sum_{i=1}^{n}\phi_{ia})^2])\Big) \\ \approx& \sum_{i=1}^{n}(\phi_{ia}-\phi_{ib})(1+\sum_{j=1}^{n}\phi_{ja}+\chi_{im})+\Big(\sum_{i=1}^{n}(\chi_{im}+1)(\phi_{ib}-\phi_{ia})+\frac{1}{2}(\sum_{i=1}^{n}\phi_{ib})^2-\frac{1}{2}(\sum_{i=1}^{n}\phi_{ia})^2\Big) \\ &-\frac{v_m}{2}\Big(\sum_{i=1}^{n}(\chi_{im}+1)(\phi_{ib}-\phi_{ia})\Big)^2 \\ =&\frac{1}{2}\Big(\sum_{i=1}^{n}(\phi_{ia}-\phi_{ib})\Big)^2-\frac{v_m}{2}\Big(\sum_{i=1}^{n}(\chi_{im}+1)(\phi_{ib}-\phi_{ia})\Big)^2. \end{aligned} \tag{114}$$

We define the effective interaction parameter as the average of the interaction parameters weighted by the difference of volume fraction of the proteins in the dense and dilute phase:

$$\chi_{\text{eff}}\equiv\sum_{i=1}^{n}\chi_{mi}\frac{\phi_{ib}-\phi_{ia}}{\sum_{j=1}^{n}(\phi_{jb}-\phi_{ja})}. \tag{115}$$

Subsequently, we obtain the zero-level curve, which is consistent with the 3-component system we discuss in section F:

$$v_m(\chi_{\text{eff}}+1)^2=1. \tag{116}$$

For $v_m>1/(\chi_{\text{eff}}+1)^2$, $Q<0$, $(\partial\phi_{1a}/\partial\bar{\phi}_m)|_{\bar{\phi}_m\to 0}<0$, so the addition of large-volume macromolecules favors phase separation, and vice versa. We remark that the above derivation equally applies to all the proteins, and the same zero-level curve applies to $(\partial\phi_{ia}/\partial\bar{\phi}_m)|_{\bar{\phi}_m\to 0}$ for $i=1\sim n$.

##### H2. The case with a major protein component

Now consider another case where a major scaffold protein component is present, and the volume fractions of all other components are much smaller than that of the major component. For simplicity, we consider a four-component system consisting of a major component 1, a minor component 2, a macromolecule 3 to be added, and solvent. We remark that in this case, the effect of adding the macromolecule on the volume fraction of the major component is the same as in the 3-component system, since the minor component's volume fraction is much smaller than the major component's.

We consider the effect of macromolecule addition to the volume fraction of the minor component. From Eq. (107), we have

$$\frac{\partial \phi_{2a}}{\partial \bar{\phi}_3} = \frac{|\mathbf{M}'|}{|\mathbf{M}|}, \tag{117}$$

where

$$\begin{aligned}
|\mathbf{M}| &= \begin{vmatrix} \mathbf{H}^a & -\mathbf{H}^b & \mathbf{0} \\ \phi_a^{\mathrm{T}}\mathbf{H}^a & -\phi_b^{\mathrm{T}}\mathbf{H}^b & 0 \\ f_a\mathbf{I_m} & (1-f_a)\mathbf{I_m} & \phi_a-\phi_b \end{vmatrix} = \begin{vmatrix} \mathbf{H}^a & -\mathbf{H}^b & \mathbf{0} \\ \mathbf{0} & (\phi_a^{\mathrm{T}}-\phi_b^{\mathrm{T}})\mathbf{H}^b & 0 \\ f_a\mathbf{I_m} & (1-f_a)\mathbf{I_m} & \phi_a-\phi_b \end{vmatrix} = \begin{vmatrix} \mathbf{0} & -\mathbf{H}^b - \frac{1-f_a}{f_a}\mathbf{H}^a & -\frac{1}{f_a}\mathbf{H}^a(\phi_a-\phi_b) \\ \mathbf{0} & (\phi_a^{\mathrm{T}}-\phi_b^{\mathrm{T}})\mathbf{H}^b & 0 \\ f_a\mathbf{I_m} & (1-f_a)\mathbf{I_m} & \phi_a-\phi_b \end{vmatrix} \\
&= |f_a\mathbf{I_m}| \cdot \begin{vmatrix} -\mathbf{H}^b - \frac{1-f_a}{f_a}\mathbf{H}^a & -\frac{1}{f_a}\mathbf{H}^a(\phi_a-\phi_b) \\ (\phi_a^{\mathrm{T}}-\phi_b^{\mathrm{T}})\mathbf{H}^b & 0 \end{vmatrix} = f_a^{n+1} \cdot \frac{(-1)^{n+1}}{f_a^{n+1}} \begin{vmatrix} f_a\mathbf{H}^b + (1-f_a)\mathbf{H}^a & \mathbf{H}^a(\phi_a-\phi_b) \\ (\phi_a^{\mathrm{T}}-\phi_b^{\mathrm{T}})\mathbf{H}^b & 0 \end{vmatrix} \\
&= (-1)^{n+1} \begin{vmatrix} \mathbf{V} & \mathbf{H}^a(\phi_a-\phi_b) \\ (\phi_a^{\mathrm{T}}-\phi_b^{\mathrm{T}})\mathbf{H}^b & 0 \end{vmatrix} = (-1)^{n+1} \begin{vmatrix} \mathbf{V} & \mathbf{H}^a(\phi_a-\phi_b) \\ \mathbf{0} & -(\phi_a^{\mathrm{T}}-\phi_b^{\mathrm{T}})\mathbf{H}^b\mathbf{V}^{-1}\mathbf{H}^a(\phi_a-\phi_b) \end{vmatrix} \\
&= (-1)^{n+1}|\mathbf{V}| \cdot \left(-(\phi_a^{\mathrm{T}}-\phi_b^{\mathrm{T}})\mathbf{H}^b\mathbf{V}^{-1}\mathbf{H}^a(\phi_a-\phi_b)\right) = (-1)^n|\mathbf{V}|(\phi_a^{\mathrm{T}}-\phi_b^{\mathrm{T}})\mathbf{H}^b\mathbf{V}^{-1}\mathbf{H}^a(\phi_a-\phi_b),
\end{aligned} \tag{118}$$

$n = 2$, and $\mathbf{V} = (1-f_a)\mathbf{H}^a + f_a\mathbf{H}^b$ is positive-definite.We note that Eq. (118) is generally valid for $n \geq 1$. Therefore, the sign of $(\partial \phi_{2a}/\partial \bar{\phi}_3)$ depends on $|\mathbf{M}'|$:

$$
\begin{aligned}
|\mathbf{M}'| &= \begin{vmatrix}
f_{11}^a & 0 & f_{13}^a & -f_{11}^b & -f_{12}^b & -f_{13}^b & 0 \\
f_{12}^a & 0 & f_{23}^a & -f_{12}^b & -f_{22}^b & -f_{23}^b & 0 \\
f_{13}^a & 0 & f_{33}^a & -f_{13}^b & -f_{23}^b & -f_{33}^b & 0 \\
\sum_{i=1}^3 \phi_{ia} f_{1i}^a & 0 & \sum_{i=1}^3 \phi_{ia} f_{3i}^a & -\sum_{i=1}^3 \phi_{ib} f_{1i}^b & -\sum_{i=1}^3 \phi_{ib} f_{2i}^b & -\sum_{i=1}^3 \phi_{ib} f_{3i}^b & 0 \\
f_a & 0 & 0 & 1-f_a & 0 & 0 & \phi_{1a}-\phi_{1b} \\
0 & 0 & 0 & 0 & 1-f_a & 0 & \phi_{2a}-\phi_{2b} \\
0 & 1 & f_a & 0 & 0 & 1-f_a & \phi_{3a}-\phi_{3b}
\end{vmatrix} \\
&= - f_a(\phi_{2a}-\phi_{2b}) \begin{vmatrix}
f_{13}^a & -f_{11}^b & -f_{12}^b & -f_{13}^b \\
f_{23}^a & -f_{12}^b & -f_{22}^b & -f_{23}^b \\
f_{33}^a & -f_{13}^b & -f_{23}^b & -f_{33}^b \\
\sum_{i=1}^3 (\phi_{ia}-\phi_{ib}) f_{3i}^a & 0 & 0 & 0
\end{vmatrix} \\
&\quad - (1-f_a)\Big((\phi_{1a}-\phi_{1b}) \begin{vmatrix}
f_{11}^a & f_{13}^a & -f_{11}^b & -f_{13}^b \\
f_{12}^a & f_{23}^a & -f_{12}^b & -f_{23}^b \\
f_{13}^a & f_{33}^a & -f_{13}^b & -f_{33}^b \\
\sum_{i=1}^3 (\phi_{ia}-\phi_{ib}) f_{1i}^a & \sum_{i=1}^3 (\phi_{ia}-\phi_{ib}) f_{3i}^a & 0 & 0
\end{vmatrix} \\
&\quad + (\phi_{2a}-\phi_{2b}) \begin{vmatrix}
f_{11}^a & f_{13}^a & -f_{12}^b & -f_{13}^b \\
f_{12}^a & f_{23}^a & -f_{22}^b & -f_{23}^b \\
f_{13}^a & f_{33}^a & -f_{23}^b & -f_{33}^b \\
\sum_{i=1}^3 (\phi_{ia}-\phi_{ib}) f_{1i}^a & \sum_{i=1}^3 (\phi_{ia}-\phi_{ib}) f_{3i}^a & 0 & 0
\end{vmatrix}\Big),
\end{aligned} \tag{119}
$$

which is a linear function of $f_a$. First we consider the case where $f_a = 1$. As $\bar{\phi}_2 \to 0$ and $\bar{\phi}_3 \to 0$, $f_{22}$ and $f_{33}$ scale as $\bar{\phi}_2^{-1}$ and $\bar{\phi}_3^{-1}$, respectively. Therefore, we can leave only the terms containing $f_{22}$ and $f_{33}$:

$$
\begin{aligned}
|\mathbf{M}'|_{f_a=1} &= -(\phi_{2a}-\phi_{2b}) \begin{vmatrix}
f_{13}^a & -f_{11}^b & -f_{12}^b & -f_{13}^b \\
f_{23}^a & -f_{12}^b & -f_{22}^b & -f_{23}^b \\
f_{33}^a & -f_{13}^b & -f_{23}^b & -f_{33}^b \\
\sum_{i=1}^3 (\phi_{ia}-\phi_{ib}) f_{3i}^a & 0 & 0 & 0
\end{vmatrix} \\
&\approx -(\phi_{2a}-\phi_{2b})\Big(\sum_{i=1}^3 (\phi_{ia}-\phi_{ib}) f_{3i}^a\Big) f_{11}^b f_{22}^b f_{33}^b \\
&\approx -\frac{f_{11}^b}{v_3\phi_{3b}} \frac{\phi_{2a}-\phi_{2b}}{v_2\phi_{2b}} \Big((\phi_{1a}-\phi_{1b}) f_{13}^a + \frac{\phi_{3a}-\phi_{3b}}{v_3\phi_{3a}}\Big).
\end{aligned} \tag{120}
$$

In the above derivation, we use $\sum_{i=1}^3 (\phi_{ia}-\phi_{ib}) f_{3i}^a \approx (\phi_{1a}-\phi_{1b}) f_{13}^a + (\phi_{3a}-\phi_{3b})/(v_3\phi_{3a})$. In the following, we will also use the approximation $\sum_{i=1}^3 (\phi_{ia}-\phi_{ib}) f_{1i}^a \approx (\phi_{1a}-\phi_{1b}) f_{11}^a$. From equilibrium of chemical potentials [Eq. (38)], we have

$$
\frac{\phi_{2a}}{\phi_{2b}} = \exp\Big(v_2\Big(\chi_{12}(\phi_{1b}-\phi_{1a}) - \ln\frac{1-\phi_{1b}}{1-\phi_{1a}}\Big)\Big), \tag{121}
$$

$$\frac{\phi_{3b}}{\phi_{3a}} = \exp\Big(-v_3\Big(\chi_{13}(\phi_{1b}-\phi_{1a}) - \ln\frac{1-\phi_{1b}}{1-\phi_{1a}}\Big)\Big). \tag{122}$$

In the biologically relevant scenario ($0 < \phi_{1a} \ll \phi_1^c < \phi_{1b} \ll 1$), we can ignore $\phi_{1a}$ and expand to the second order of $\phi_{1b}$ to get

$$\frac{\phi_{2a}}{\phi_{2b}} \approx 1 + v_2\Big(\Delta\chi_{12}\phi_{1b} + \frac{1}{2}\phi_{1b}^2\Big) + \frac{1}{2}v_2^2\Delta\chi_{12}^2\phi_{1b}^2, \tag{123}$$

$$\frac{\phi_{3b}}{\phi_{3a}} \approx 1 - v_3\Big(\Delta\chi_{13}\phi_{1b} + \frac{1}{2}\phi_{1b}^2\Big) + \frac{1}{2}v_3^2\Delta\chi_{13}^2\phi_{1b}^2. \tag{124}$$

Here, $\Delta\chi_{12} \equiv \chi_{12}+1$ and $\Delta\chi_{13} \equiv \chi_{13}+1$. We also have $f_{13}^a = 1/(1-\phi_{1a}-\phi_{2a}-\phi_{3a}) + \chi_{13} \approx \Delta\chi_{13}$. Plugging these approximations into Eq. (120), we get

$$|\mathbf{M}'|_{f_a=1} \approx \frac{f_{11}^b}{2v_3\phi_{3b}}\Delta\chi_{12}(v_3\Delta\chi_{13}^2 - 1)\phi_{1b}^3. \tag{125}$$

Then we consider the case where $f_a = 0$. Leaving only the terms containing $f_{22}$ and $f_{33}$, we get

$$\begin{aligned}
|\mathbf{M}'|_{f_a=0} =& \frac{1}{v_3\phi_{3b}}\Big\{(\phi_{1a}-\phi_{1b})\Big[(\phi_{1a}-\phi_{1b})f_{11}^a\frac{\phi_{3b}}{\phi_{3a}}(f_{11}^b f_{23}^b - f_{12}^b f_{13}^b) + (\phi_{1a}-\phi_{1b})f_{11}^a(f_{13}^a f_{12}^b - f_{11}^b f_{23}^a)\\
&-\Big((\phi_{1a}-\phi_{1b})f_{13}^a + \frac{\phi_{3a}-\phi_{3b}}{v_3\phi_{3a}}\Big)(f_{11}^a f_{12}^b - f_{12}^a f_{11}^b)\Big]\\
&-\frac{\phi_{2a}-\phi_{2b}}{v_2\phi_{2b}}\Big[(\phi_{1a}-\phi_{1b})f_{11}^a f_{13}^b\frac{\phi_{3b}}{\phi_{3a}} + f_{11}^a\Big((\phi_{1a}-\phi_{1b})f_{13}^a + \frac{\phi_{3a}-\phi_{3b}}{v_3\phi_{3a}}\Big) - (\phi_{1a}-\phi_{1b})f_{13}^a f_{11}^a\Big]\Big\}.
\end{aligned} \tag{126}$$

Notice that since $\phi_{1a} \ll \phi_{1b}$, $f_{11}^a \gg f_{11}^b$. Therefore, we can ignore the term without $f_{11}^a$. Plugging in the following approximations (remember that in biologically relevant scenario, $\phi_{1b} \approx 3\Delta\chi_{11}/2, \Delta\chi_{11} \equiv -\chi_{11} - 1$):

$$f_{11}^b = \frac{1}{v_1\phi_{1b}} + \frac{1}{1-\phi_{1b}-\phi_{2b}-\phi_{3b}} + \chi_{11} \approx \chi_{11} + 1 + \phi_{1b} = \phi_{1b} - \Delta\chi_{11} \approx \frac{1}{3}\phi_{1b}, \tag{127}$$

$$f_{12}^b = \frac{1}{1-\phi_{1b}-\phi_{2b}-\phi_{3b}} + \chi_{12} \approx \Delta\chi_{12} + \phi_{1b}, \quad f_{13}^b \approx \Delta\chi_{13} + \phi_{1b}, \quad f_{23}^b \approx \Delta\chi_{23} + \phi_{1b}, \tag{128}$$

$$f_{12}^a = \frac{1}{1-\phi_{1a}-\phi_{2a}-\phi_{3a}} + \chi_{12} \approx \Delta\chi_{12}, \quad f_{13}^a \approx \Delta\chi_{13}, \quad f_{23}^a \approx \Delta\chi_{23}, \tag{129}$$

and the approximations of $\phi_{2a}/\phi_{2b}$ and $\phi_{3b}/\phi_{3a}$, we expand Eq. (126) to the fourth order of $\phi_{1b}$:

$$|\mathbf{M}'|_{f_a=0} \approx \frac{f_{11}^a}{v_3\phi_{3b}}\Big(\frac{1}{12} + \frac{1}{4}v_3\Delta\chi_{13}^2 - \frac{1}{3}v_3\Delta\chi_{13}\Delta\chi_{23} + \frac{1}{4}(1 - v_3\Delta\chi_{13}^2)v_2\Delta\chi_{12}^2\Big)\phi_{1b}^4. \tag{130}$$

Finally, we have

$$|\mathbf{M}'| = f_a|\mathbf{M}'|_{f_a=1} + f_b|\mathbf{M}'|_{f_a=0}. \tag{131}$$

We note that $|\mathbf{M}'|_{f_a=0} \propto f_{11}^a \propto \phi_{1a}^{-1}$, which is much larger than $|\mathbf{M}'|_{f_a=1}$ in the biologically relevant scenario; therefore, we can neglect $|\mathbf{M}'|_{f_a=1}$ when $f_b$ is not too close to 0. Under this condition, we have

$$\frac{\partial\phi_{2a}}{\partial\bar{\phi}_3} \propto \frac{1}{12} + \frac{1}{4}v_3\Delta\chi_{13}^2 - \frac{1}{3}v_3\Delta\chi_{13}\Delta\chi_{23} + \frac{1}{4}(1 - v_3\Delta\chi_{13}^2)v_2\Delta\chi_{12}^2. \tag{132}$$

Fig. S4 shows the sign of $\partial\phi_{2a}/\partial\bar{\phi}_3$ under different conditions. Dots are calculated directly from Eq. (117). Red dots represent positive $\partial\phi_{2a}/\partial\bar{\phi}_3$, and blue dots represent negative $\partial\phi_{2a}/\partial\bar{\phi}_3$. The zero-level curves are calculated from the approximated result in Eq. (132).

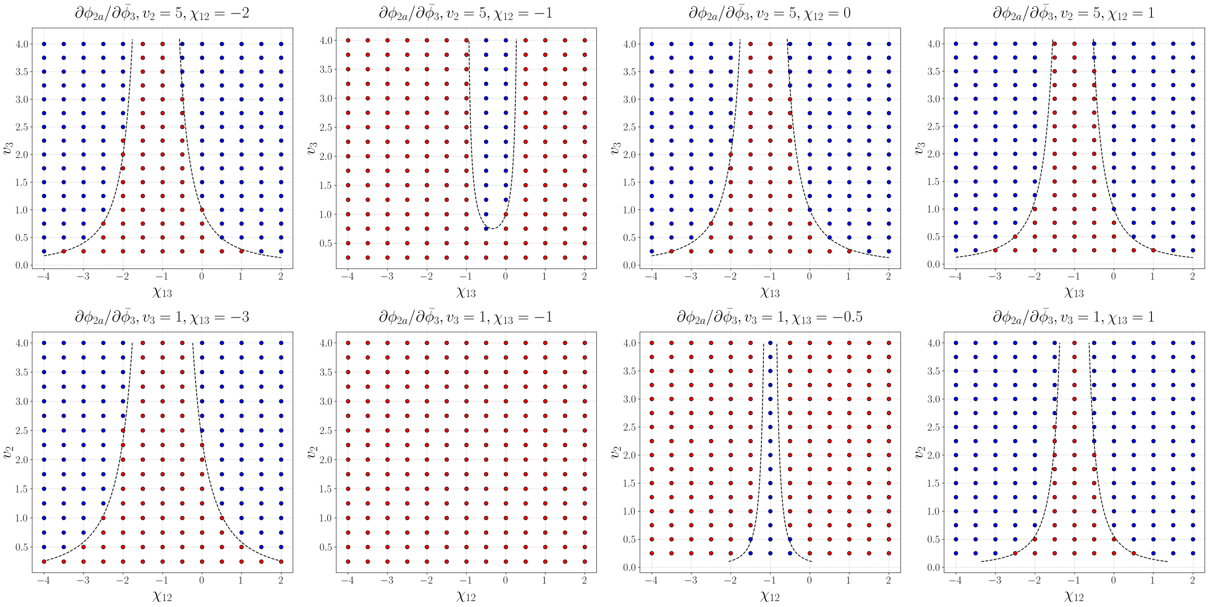


FIG. S4. Sign of $\partial\phi_{2a}/\partial\bar{\phi}_3$. Each subfigure in the first row is calculated at fixed $v_2$ and $\chi_{12}$; each subfigure in the second row is calculated at fixed $v_3$ and $\chi_{13}$. Red dots represent positive values and blue dots represent negative values. In all subfigures, $v_1 = 100000$,$\chi_{11} = -1.02$, $f_b = 0.2$, $\chi_{22} = \chi_{23} = \chi_{33} = 0$.

### I. DISCUSSION ON THE NON-MEAN-FIELD MODEL

We consider phase separation of a protein in poor solvent under the following condition:

$$v_1 \gg 1, \frac{1}{\sqrt{v_1}} \ll \Delta\chi_{11} \ll 1. \tag{133}$$

The dense phase is well described by mean-field theory:

$$f^b = \frac{1}{v_1}\phi_{1b}\ln\phi_{1b} + (1-\phi_{1b})\ln(1-\phi_{1b}) + \frac{1}{2}\chi_{11}\phi_{1b}^2. \tag{134}$$

However, in the dilute phase, the protein chains exist as isolated globules. Therefore, the mean-field theory does not hold for the dilute phase.

#### I1.Chemical potential equilibrium

We remark that the volume fraction of the protein inside the globule is the same as that in the dense phase [1]:

$$\phi_{\text{in}} = \frac{v_1}{\frac{4}{3}\pi R_{gl}^3} \approx \phi_{1b}. \tag{135}$$

Therefore, the part of the chemical potential generated by bulk interaction is the same between the globule and the dense phase. Therefore, to compute the volume fraction in the dilute phase, we only need to take into account the surface energy of the globule and the translational entropy of the globule.

Next we consider the surface energy of a globule using blob theory. The size of a thermal blob scales with the excluded volume $v$, which is proportional to $\Delta\chi_{11}$ [1]:

$$\xi \approx \frac{b^4}{|v|} \approx \Delta\chi_{11}^{-1}. \tag{136}$$

We omit the monomer size b as it is set to be the length unit (and the monomer volume is the volume unit).

The surface tension $\gamma$ arises from the energy penalty of blobs at the surface interacting with the solvent. The energy cost is approximately $k_BT$ per blob area $\xi^2$ (here we omit energy unit $k_BT$):

$$\gamma \approx \frac{k_BT}{\xi^2} \approx (\Delta\chi_{11})^2. \tag{137}$$

The total surface energy of a globule with radius $R_{gl}$ is

$$E_{\text{surface}} = 4\pi R_{gl}^2\gamma \approx R_{gl}^2(\Delta\chi_{11})^2. \tag{138}$$

Density of globule in the dilute phase is $\phi_{1a}/v_1$. Therefore,

$$e_{\text{surface}} = \frac{\phi_{1a}}{v_1}E_{surface} \approx \frac{R_{gl}^2(\Delta\chi_{11})^2}{v_1}\phi_{1a}, \tag{139}$$

and the chemical potential of the protein in the dilute phase generated by surface energy can be expressed as

$$\mu_{\text{surface}} = \frac{\partial e_{\text{surface}}}{\partial\phi_{1a}} \approx \frac{R_{gl}^2(\Delta\chi_{11})^2}{v_1}. \tag{140}$$

The translational entropy is $-(k_B\phi_1\ln\phi_1)/v_1$ in both dilute and dense phase, and it contributes $(\ln\phi_1+1)/v_1$ to the chemical potential. Because the difference of contribution from translational entropy must equal the contribution of surface energy, we obtain

$$\frac{1}{v_1}\ln\phi_{1b} - \frac{1}{v_1}\ln\phi_{1a} \approx \frac{R_{gl}^2(\Delta\chi_{11})^2}{v_1}. \tag{141}$$

##### I2. Osmotic pressure

From Eq. (141) we deduce that $\phi_{1a}$ is exponentially smaller than $\phi_{1b}$. Therefore, the osmotic pressure of the dilute phase is negligible. We have

$$0 = \Pi^a = \Pi^b = \Big(\frac{1}{v_1}-1\Big)\phi_{1b} - \ln(1-\phi_{1b}) + \frac{1}{2}\chi_{11}\phi_{1b}^2, \tag{142}$$

which is the same as what we derived in section F. Therefore, we still have

$$\phi_{1b} \approx \frac{3}{2}\Delta\chi_{11}. \tag{143}$$

Plugging into Eq. (135), we get

$$R_{gl} \approx \Big(\frac{3v_1}{4\pi\phi_{1b}}\Big)^{1/3} \approx \Big(\frac{v_1}{2\pi\Delta\chi_{11}}\Big)^{1/3}. \tag{144}$$

Substituting the expression of $R_{gl}$ into Eq. (141), we get

$$\phi_{1a} \approx \phi_{1b}\exp(-R_{gl}^2(\Delta\chi_{11})^2) \approx \frac{3}{2}\Delta\chi_{11}exp(-C\Delta\chi_{11}^{4/3}v_1^{2/3}), \tag{145}$$

where $C$ is a constant.

##### I3. Calculation of $(\partial\phi_{1a}/\partial\bar{\phi}_2)|_{\bar{\phi}_2\to 0}$ and $(\partial\phi_{1b}/\partial\bar{\phi}_2)|_{\bar{\phi}_2\to 0}$

We remark that all the derivations for this non-mean-field case are essentially the same as what we do for Flory-Huggins theory, except for the expression of $\phi_{1a}$. For example, in Eq. (95), one only needs to replace $\phi_{1a}$ by the non-mean-field expression. Therefore,

$$\left.\frac{\partial\phi_{1b}}{\partial\bar{\phi}_2}\right|_{\bar{\phi}_2\to 0} \approx \frac{3}{2}(v_2\Delta\chi_{12}^2-1), \tag{146}$$

$$\left.\frac{\partial \phi_{1a}}{\partial \bar{\phi}_2}\right|_{\bar{\phi}_2 \to 0} \approx \frac{1}{2}(1 - v_2 \Delta\chi_{12}^2)\phi_{1b}^2 \frac{v_1 \phi_{1a}}{\phi_{1b}} \\ \approx \frac{9}{8}(1 - v_2 \Delta\chi_{12}^2) v_1 \Delta\chi_{11}^2 exp(-C\Delta\chi_{11}^{4/3} v_1^{2/3}). \tag{147}$$

The expression for the zero-level curves does not change. We note that our analysis of the non-mean-field effects is preliminary here, as we neglect some potentially important factors (e.g., the non-mean-field effects on $\phi_2$), and we expect more systematic studies of the non-mean-field models in the future.

[1] Michael Rubinstein and Ralph H Colby. *Polymer physics.* Oxford university press, 2003.
